# Neural Population Models for EEG: From Canonical Models to Alternative Model Structures

**DOI:** 10.64898/2026.04.10.717643

**Authors:** Nina Omejc, Sabin Roman, Ljupčo Todorovski, Sašo Džeroski

**Affiliations:** Department of Knowledge Technologies, Jožef Stefan Institute, Ljubljana, Slovenia; Jožef Stefan International Postgraduate School, Ljubljana, Slovenia; Department of Mathematics, Faculty of Mathematics and Physics, University of Ljubljana, Slovenia

## Abstract

Neural population models are widely used to interpret electroencephalography (EEG), yet their relationships remain far less systematically understood than those among single-neuron models. More fundamentally, it remains unclear whether EEG can support a uniquely plausible population-level mechanism, or whether multiple structurally distinct models can explain the data equally well. To address this question, we combine comparative analysis of canonical model families with grammar-based generation of new candidate architectures. We assembled 17 canonical neural mass and phenomenological models and embedded them in a shared structural space. From their common processes, we defined a probabilistic grammar over interpretable dynamical components and developed ENEEGMA (Exploring Neural EEG Model Architectures), a Julia-based framework for grammar-based model generation, simulation, and parameter optimization, to generate additional candidate models. We then assessed both canonical and generated models by fitting them to EEG independent-component spectra from four datasets for each condition: resting state and steady-state visual evoked potentials. Canonical models formed six structural clusters. Across conditions, compact low-dimensional polynomial oscillators performed best overall, with Montbrió–Pazó–Roxin, FitzHugh–Nagumo, and Stuart–Landau models offering the best balance of fit quality, stability, and simplicity. Grammar-based exploration further showed that the space of viable EEG node models extends beyond canonical formulations: even a restricted search over 1,000 generated models produced compact alternatives competitive with nearly all canonical families and achieving the strongest cluster-level SSVEP fits. Together, these findings suggest that EEG spectra constrain plausible neural population mechanisms without uniquely determining them. Beyond this, grammar-based model exploration provides a principled, data-driven framework for EEG-constrained model discovery.

**Author summary:** Electroencephalography (EEG) lets us measure brain activity non-invasively, but the signals are indirect, so we rely on mathematical models to explain how neural populations generate them. Many such models exist, yet it is unclear whether standard models cover the full range of plausible explanations for EEG data, or whether several very different models can explain the same signal equally well. In this study, we compared a broad set of established neural population models and then used a grammar-based equation discovery framework to automatically generate new candidate models from interpretable building blocks. We found that simple low-dimensional oscillator models often matched EEG spectra better than more complex canonical models. We also found that newly generated models could perform nearly as well as, and sometimes better than, established ones, especially for stimulus-driven responses. These results suggest that EEG spectra alone may not be enough to identify a unique underlying neural mechanism. More broadly, our work shows how automated, biologically informed model generation can help to compare, understand, expand, and test the space of candidate neural population models.

## Introduction

In humans, non-invasive recording techniques provide a major source of information about population-level brain dynamics. In particular, electroencephalography (EEG) offers millisecond temporal resolution. However, it provides only indirect and spatially blurred access to the underlying neural population dynamics, which are additionally nonlinear, high-dimensional, and intrinsically noisy [1–3]. Interpreting these signals, therefore, relies on modelling approaches that link neuronal mechanisms to macroscopic EEG observables. These approaches range from classical, biophysically grounded dynamical systems [4–7] to more recent, prediction-oriented data-driven neural networks [8, 9]. Here, we focus on biophysically grounded dynamical systems that describe the collective dynamics underlying EEG signals.

### Modelling EEG

Mathematical models of neural populations span a spectrum, from detailed single-neuron and spiking-network models [10], through population-density and neural mass formulations [4, 11, 12], to neural-field models [6, 7] and highly reduced phenomenological dynamical systems [13, 14]. In this work, we focus on the low-dimensional end of this spectrum, specifically neural mass and phenomenological models, which can be studied within a shared framework of low-dimensional dynamical systems, including ordinary, stochastic, and delay differential equations. These modelling regimes exhibit a particularly broad diversity of formulations and comparatively fewer standardized reference models than detailed single-neuron and spiking-network models. This combination creates both a practical need and a growing curiosity about the modelling space, highlighting the value of systematic analysis and principled exploration.

Some of the earliest neural mass models were introduced by Lopes da Silva [15], Wilson and Cowan [16], and Jansen and Rit [17]. These models describe the collective dynamics of interacting neuronal populations under the assumption that each population is spatially lumped and internally homogeneous. Their state variables represent population-averaged quantities such as mean membrane potential, mean firing rate, or synaptic current, resulting in low-dimensional dynamical systems. Neural mass models are typically formulated as systems of ordinary differential equations of the form 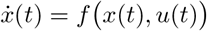, often augmented with stochastic fluctuations or transmission delays to capture variability and finite propagation effects [4, 12]. Here, *x*(*t*) denotes the vector of population state variables, while *u*(*t*) represents external or afferent input to the population. Representative applications include methodological and theoretical studies of seizure, resting-state, and network dynamics [18–25], as well as clinical and translational modeling in disease-related settings [26, 27].

Phenomenological models, by contrast, often draw on concepts and mathematical structures originating in physics and dynamical systems theory [13]. They have been adapted to capture essential macroscopic dynamical features of neural activity without explicitly modelling underlying biophysical mechanisms. Despite their reduced dimensionality, such models have proven valuable for describing characteristic macroscopic neural dynamics [28], including oscillations, relaxation cycles, synchronization phenomena, and pathological transitions such as seizures [29]. Owing to their balance between mathematical simplicity, computational efficiency, and interpretability at the population level, phenomenological models are used alongside neural mass models in contemporary EEG research [28, 30–33].

The literature offers a growing catalogue of candidate neural population models [4, 12, 34]. Software ecosystems such as The Virtual Brain (TVB) [35–37], Brain Dynamics Toolbox [38], PyRates [39, 40], and Dynamic Causal Modelling (DCM) [41, 42] have made these models widely accessible, providing powerful frameworks for simulating and fitting predefined model structures. Despite the availability of these tools, several conceptual and methodological issues remain insufficiently addressed. In particular, although many neural mass and phenomenological models have been proposed, there is still limited general guidance on how canonical formulations differ in their structural assumptions, dynamical behaviour, and empirical adequacy for specific EEG datasets or scientific questions. Comparative analyses are often model-specific, and a systematic organization of the neural population model space, analogous to that available for single-neuron models, remains largely absent. As a result, it is unclear whether the current collection of canonical models adequately spans the space of plausible population-level dynamics, or whether alternative model structures, consistent with known neurophysiological constraints, could provide equally good or improved descriptions of EEG data.

Addressing these challenges requires a framework that can systematically analyze existing models while also exploring novel, biologically plausible alternatives. In practice, such exploration is difficult to carry out manually, given the size and combinatorial diversity of the candidate model space. It therefore depends on a representation of neural population models that is explicit, compositional, and suited to algorithmic search. To this end, we adopt a grammar-based equation discovery approach that probabilistically combines biologically interpretable building blocks to generate and formalize neural population models.

### Equation discovery via probabilistic grammars

From a dynamical-systems perspective, modelling an observed signal is an inverse problem: given an observed trajectory **y**(*t*), we assume an underlying dynamical system of the form 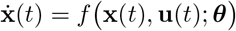, where the observed signal is related to the latent state via an observation mapping **y**(*t*) = *g*(**x**(*t*)) + ***ε***(*t*), where ***ε***(*t*) denotes measurement noise. Classical system identification assumes a known functional form *f* and estimates only the parameters ***θ***. In contrast, equation discovery (also called symbolic regression) infers both the structure of *f* and its parameters from data [43]. In simple terms, equation discovery lets a computer automatically generate and test many candidate equations or model structures, often tens of thousands or more, to identify those that best reproduce the observed data. A variety of methods have been proposed, including early equation-discovery systems such as LAGRAMGE [44, 45], sparse regression–based approaches such as SINDy [46, 47], probabilistic grammar-based frameworks such as ProGED [48, 49], and neural approaches such as DSO [50] and HVAE [51].

In neuroscience, equation-discovery approaches remain relatively unexplored. One recent study used SINDy to recover simulated low-dimensional stochastic decision dynamics in perceptual decision-making tasks, illustrating the promise of such methods in computational neuroscience [52]. To our knowledge, this is the first study to apply equation-discovery methods to identify neural population dynamics directly from EEG data. A key challenge in doing so is that searching over arbitrary analytic expressions is difficult: the space of candidate models is vast, and generic algebraic primitives encode little domain-specific knowledge about neural dynamics. To address this challenge, we incorporate domain knowledge by formalizing the search process using a probabilistic grammar, as in ProGED [48, 49], that specifies how high-level components of population models are expanded into concrete mathematical forms. A probabilistic grammar can be thought of as a set of weighted assembly rules: it tells the computer which building blocks can be combined and which combinations are more plausible. This biases equation discovery toward simpler, more interpretable, and biologically meaningful structures.

Motivated by these considerations, we developed ENEEGMA (Exploring Neural EEG Model Architectures). This framework uses a domain-informed, probabilistic grammar to generate, simulate, and evaluate neural population models for EEG data. This framework enables systematic exploration of established formulations as well as a broader model space, including combinations not previously proposed. Using ENEEGMA, we address two central questions: first, how canonical neural mass and phenomenological models compare in terms of structure and empirical performance; and second, whether systematic exploration can reveal alternative, neurophysiologically plausible architectures with comparable or improved adequacy for EEG data.

## Materials and methods

We first introduce the established, canonical neural population models and describe the structural and empirical analyses used to compare them. We then detail the construction of the probabilistic grammar, the generation of novel models, and their analysis.

### Canonical neural population models

We collected a set of canonical neural population models from The Virtual Brain (TVB) [35, 37], the Brain Dynamics Toolbox [38], PyRates [39], Dynamic Causal Modeling (DCM) [41], and recent comprehensive reviews [34, 53]. In Table 1, we list seventeen canonical models that were analysed, compared, and later also used as domain knowledge to construct the grammar.

**Table 1.** Canonical neural population models and their corresponding dynamical grammar components. Each row lists one reference model, its properties, and its coarse-grained structural and dynamical features as represented in the grammar. The columns report: *N*_pop_, the number of interacting neural populations; *N*_var/pop_, the number of state variables per population; *N*_par_, the number of model parameters in the grammar-generated model; *Input/Output processes*, the local transformation used to map inputs to population dynamics and, where applicable, the output operator; *Conn. funcs*, the functional forms used for inter- or intra-population connectivity; and *Conn. motif*, the characteristic connectivity pattern among populations. Model abbreviations are: WC, Wilson–Cowan; ARM, Alpha Rhythm Model; JR, Jansen–Rit; W, Wendling; MDF, Moran–David–Friston; LW, Liley–Wright (reduced); RRWC, Robinson–Rennie–Wright Cortex; RRWT, Robinson–Rennie–Wright Thalamus; WW, Wong–Wang; WWR, reduced Wong–Wang; LB, Larter–Breakspear; HO, Harmonic Oscillator; MPR, Montbrió–Pazó–Roxin; FHN, FitzHugh–Nagumo; VDP, Van der Pol; SL, Stuart–Landau; and DO, Duffing oscillator. Additional abbreviations are: sat. sig., saturating sigmoid; base. sig., base sigmoid; mem. integrator, membrane integrator.

| Model | $N_{\text{pop}}$ | $N_{\text{var/pop}}$ | $N_{\text{par}}$ | Input/Output processes | Conn. funcs | Conn. motif |
| --- | --- | --- | --- | --- | --- | --- |
| WC | 2 | 1 | 14 | exponential / direct | base. sig. | full |
| ARM | 2 | 2 | 10 | second-order / direct | linear | ring |
| JR | 3 | 2 | 25 | second-order / direct | sat. sig. | star |
| W | 5 | 2 | 57 | second-order / direct | sat. sig.; linear | star-tail |
| MDF | 3 | 2–5 | 35 | second-order / difference | base. sig.; linear | multi-cycle |
| LW | 2 | 5 | 41 | second-order / mem. integrator | sat. sig.; linear | chain-loop |
| RRWC | 2 | 4 | 24 | second-order / spatial gradient | sat. sig.; linear | full |
| RRWT | 2 | 2 | 16 | second-order / direct | sat. sig.; linear | ring |
| WW | 2 | 1 | 16 | gating kinetics / direct | rectifier | full |
| WWR | 1 | 1 | 7 | gating kinetics / direct | rectifier | full |
| LB | 1 | 3 | 31 | voltage-dependent / direct | linear | null |
| HO | 1 | 2 | 3 | second-order / direct | linear | null |
| MPR | 1 | 2 | 9 | polynomial / direct | linear | null |
| FHN | 1 | 2 | 10 | polynomial / direct | linear | null |
| VDP | 1 | 2 | 9 | polynomial / direct | linear | null |
| SL | 1 | 2 | 13 | polynomial / direct | linear | null |
| DO | 1 | 2 | 9 | polynomial / direct | linear | null |

Broadly, the surveyed canonical models can be roughly divided into two groups, neural mass models (NMMs) and phenomenological models. In the neural mass models category, we include Wilson–Cowan (WC) [16, 54], the Alpha Rhythm Model (ARM) [11, 15], Jansen–Rit (JR) [17], Wendling (W) [55], Moran–David–Friston (MDF) [41, 53, 56, 57], Liley–Wright (LW) [58], the Robinson–Rennie–Wright (RRW) thalamo-cortical model [59, 60], Wong–Wang (WW) models as implemented in TVB [19, 61, 62], and Larter–Breakspear (LB) [63, 64]. Several of these models were not originally formulated as ordinary differential equations, but rather as spatially extended (e.g., neural field models), and are considered here in their reduced, lumped ODE or neural mass formulations. From the set of phenomenological models, we include the harmonic oscillator (HO), FitzHugh–Nagumo (FHN) [65, 66], Van der Pol (VDP), Stuart–Landau (SL), and Duffing oscillator (DO) [13]. We also include the Montbrió–Pazó–Roxin (MPR) model [67], which is a rigorously derived mean-field reduction of spiking neuronal networks and is considered here as a low-dimensional phenomenological population model. A mathematical formulation of all models is provided in Supplementary Material A.

### Structural analysis of canonical models

We performed a structural analysis of canonical neural population models based solely on their mathematical form. Each model is represented as a system of differential equations, from which symbolic expressions of the right-hand sides are extracted. This allowed us to characterize similarities and differences between models at the level of their equations alone, independently of parameter choices and empirical data.

To compare models structurally, we combined two complementary distance measures with equal weights. The first is a syntactic edit distance between symbolic equations [68], which captures fine-grained differences in algebraic structure. It is defined as the minimum number of insertion, deletion, and substitution operations required to transform one symbolic expression into another, and is normalized to the unit interval using bounds estimated from a large reference set of grammar-sampled models. The second measure captures similarity in equation content by comparing the frequencies of operators and functions, irrespective of their ordering. Specifically, it is computed as the cosine distance between feature vectors encoding the counts of operators and functions present in the respective equations [69].

For the canonical models, we computed pairwise structural distances to construct a distance matrix, which served as the basis for clustering. Clustering was performed using agglomerative hierarchical clustering with optimal leaf ordering [70], as implemented in the Julia package Clustering.jl. This procedure enabled the identification of structurally related model families without imposing prior assumptions about model classes.

Additionally, model complexity was quantified as the total number of state equations, reflecting the dimensionality of the underlying dynamical system and serving as a proxy for structural richness and implementation cost.

### Empirical evaluation of canonical models

We performed an empirical evaluation of canonical neural population models based on their observable dynamics, assessing how well they reproduce the spectral characteristics of EEG data. This enabled comparison across models with widely differing internal structure, dimensionality, and parameterization. Rather than focusing on intrinsic dynamical properties such as fixed points, stability, or bifurcation structure, this analysis evaluates models at the level of their measurable output, enabling direct comparison across heterogeneous model classes.

#### EEG data

Recorded EEG data were obtained from the dataset of Lee et al. [71], accessed via the MOABB framework [72]. This benchmark was selected because it includes both resting-state (RS) recordings and a steady-state visual evoked potential (SSVEP) paradigm, which yield well-characterized and distinct spectral signatures. For each condition, we selected four representative independent components, one from each dataset. For the SSVEP condition, we selected a visual stimulation frequency of 6.67Hz. Detailed preprocessing of the EEG data, including independent component analysis, is described in Supplementary Material B.

#### Signal representation

Both empirical data and simulated model outputs were transformed into power spectral density (PSD) estimates using identical spectral estimation settings to ensure comparability. Using Welch’s method, signals were segmented into overlapping windows of length *n*_perseg_ = 1024 samples with 50% overlap and a Hann window. Spectra were computed over the frequency range 1–45 Hz, averaged across segments, and analyzed on a logarithmic scale (log-power spectrum; hereafter log-PSD). To reduce variability arising from noise, model spectral estimates were additionally averaged across three independent noise realizations. The resulting log-PSD provides a robust, phase-invariant summary of neural dynamics and allows direct comparison between models whose state variables have different interpretations or units.

#### External drive

All models were driven by externally applied input signals constructed to reflect the experimental condition. For RS simulations, models received white noise, providing a common source of background fluctuations across models, including otherwise deterministic formulations. For SSVEP simulations, we used a periodic sinusoid driving component at the stimulation frequency of 6.67 Hz to enable stimulus-locked responses. Note that the input consisted of a pure sinusoid without harmonic components.

#### Objective function

Model fitting was guided by a composite spectral objective defined in log-PSD space.

We define a masked mean absolute error (MAE) as

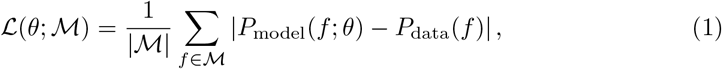

where *P*_data_(*f*) and *P*_model_(*f* ; *θ*) denote the empirical and simulated log-PSD evaluated over the frequency grid ℱ, and ℳ ⊆ ℱ specifies the frequency bins included in the loss.

The total loss for the resting state is

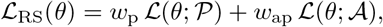

where 𝒫 contains detected spectral peaks and 𝒜 = ℱ\ 𝒫 captures broadband aperiodic structure. We fixed *w*_p_ to 1 and treated *w*_ap_ as a hyperparameter, varying it between 0 and 1.

In SSVEP, for stimulation frequency *f*_0_, let ℋ denote frequency bins around *f*_0_ and its first *H* harmonics. The harmonic contribution is

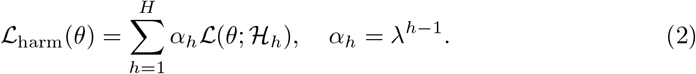

The total SSVEP loss becomes

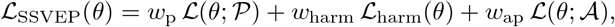

where 𝒜 = ℱ\ (𝒫 = ℋ). We fixed *w*_harm_ to 1 and treated *w*_p_ and *w*_ap_ as coupled hyperparameters, varying together between 0 and 1.

#### Parameter estimation

Parameter estimation was formulated as a model-agnostic optimization problem in which tunable parameters and unknown initial conditions were adjusted to minimize the spectral objective defined above. Optimization was performed using the Covariance Matrix Adaptation Evolution Strategy (CMA-ES) [73], a derivative-free method suited to nonconvex problems.

Ordinary differential equations were integrated using an adaptive solver that switches between stiff (Rodas5) and non-stiff (Tsit5) regimes, while stochastic systems were simulated using the Euler–Maruyama method [74]. Additive measurement noise, with amplitude estimated from empirical data, was included in simulations.

To ensure numerical stability and comparable scaling across heterogeneous parameter sets, parameters were internally reparameterized into a shared dimensionless optimization space.

Hyperparameters for canonical models were selected using an exhaustive grid search on the first dataset, varying four hyperparameters. First, for the RS condition, we varied *w*_*ap*_ (0.5, 0.75, 1.0) while fixing the task-relevant spectral weights to 1, whereas for the SSVEP condition, we fixed the task-relevant harmonic weight *w*_*harm*_ to 1 and varied *w*_*p*_ = *w*_*ap*_ jointly over (0.5, 0.75, 1.0). Second, parameter defaults were initialized from model-specific literature values where available. We varied the width of the optimization bounds around these default values using two alternative settings: a narrower range from 0.25 to 4 times the default value, and a broader range from 0.125 to 8 times the default value. For parameters with zero default value, symmetric fallback intervals of [−5, 5] and [−10, 10] were used, respectively. Third, we varied the optimizer initial step size *σ*_0_ (2, 8), and fourth, the optimizer population size (100, 150). This yielded 24 configurations per model. During hyperparameter tuning, each configuration was evaluated in five independent optimization runs initialized from different starting parameter values, for a total of 120 runs per model. The best-performing setting was then fixed and used to fit the remaining three datasets for each condition.

#### Evaluation

For the final evaluation, the best-fitting parameter set for each model was fixed, and the model was re-simulated five times with different initial conditions and noise realizations, yielding five evaluation runs per dataset. This allowed us to assess the robustness of the fitted solution.

Final model comparison and statistical analysis were based on the integrated absolute error (IAE) over the fitting range ℱ,

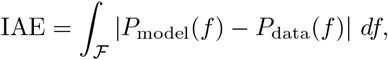

computed numerically by trapezoidal integration. This avoids dependence on the optimization weighting scheme and provides a directly interpretable, distribution-robust measure of spectral discrepancy that does not assume Gaussian residual structure. For SSVEP data, we additionally computed a harmonic-restricted IAE (h-IAE), defined analogously to IAE, but with the integration domain restricted from ℱ to the stimulus-locked harmonic regions ℋ ⊂ ℱ. This additional metric ensures that model ranking is not mainly driven by the accurate fit of broadband background aperiodic structure, but reflects fidelity to the evoked harmonic response.

To compare models at the group level, performance summaries were aggregated across datasets. Because only four independent datasets per condition were included, we used a nonparametric Bayesian bootstrap [75]. In simple terms, this method repeatedly reweights the observed datasets, recomputes the group-level summaries, and ranks the models, thereby quantifying uncertainty in model comparison without relying on strong distributional assumptions. From these posterior samples, we estimated pairwise posterior probabilities of model superiority, posterior probabilities of being the best-performing model, and Bayesian expected ranks.

### Probabilistic grammar and grammar-derived candidate models

To enable systematic exploration beyond existing neural population models, we developed ENEEGMA. This unified framework integrates grammar-based model generation with the simulation, optimization, and evaluation pipeline described above and graphically in Fig. 1. Candidate node-level models are generated by sampling from a probabilistic grammar that encodes construction rules derived from canonical neural population models and specifies how these components are combined into complete systems of differential equations. While ENEEGMA is designed to embed node models within atlas-registered whole-brain networks that incorporate inter-regional coupling and transmission delays, this study restricts the analysis to single-node models. Comprehensive network-level modelling and evaluation are beyond the scope of the present work.

**Fig 1.**
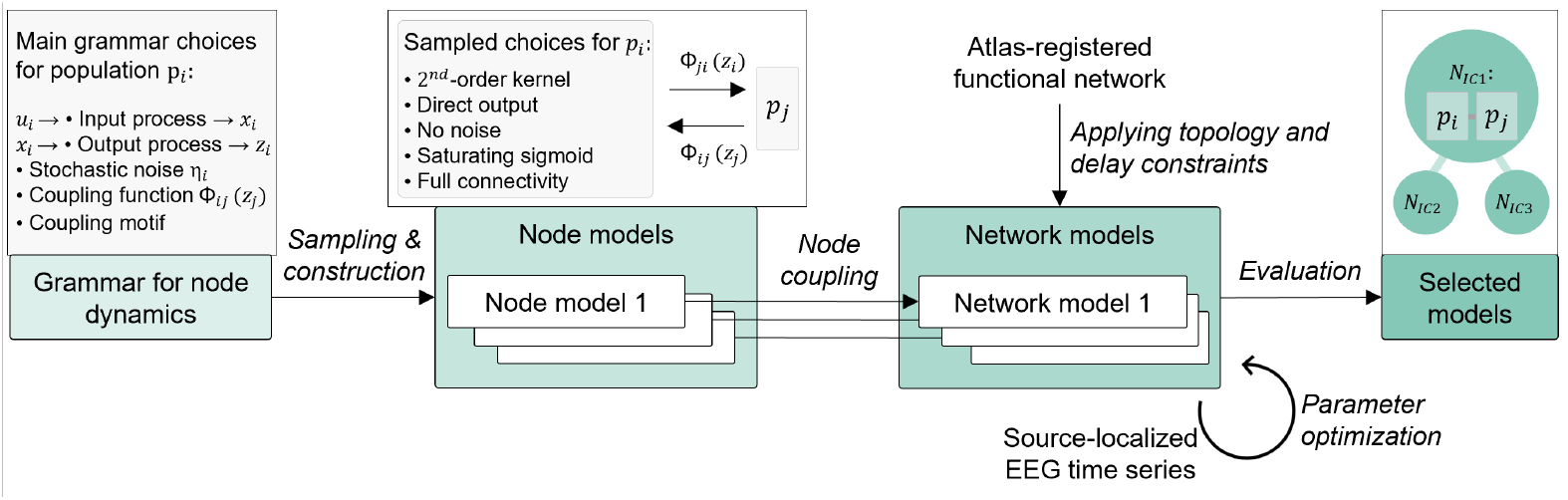
High-level overview of the ENEEGMA pipeline. A probabilistic grammar defines admissible node dynamics in terms of interacting populations, each specified by a selected input process, output process, optional stochasticity, coupling functions, and a connectivity motif. Sampling and construction from this grammar yield candidate node models. These node models are then embedded into atlas-registered functional networks under topology and delay constraints to form network models. The resulting network models are evaluated against source-localized EEG spectral features through parameter optimization, and the best-performing models are selected. Green boxes denote the main components of the ENEEGMA pipeline, italic labels on the arrows indicate the main workflow stages, and gray/white boxes illustrate example components at each stage.

By examining the canonical models, we observe that they can be decomposed into a small number of interacting populations, typically ranging from two to five. We further decompose the dynamics of each population into four main components: (i) *Input processes*, which integrate presynaptic firing rates into internal drive variables; (ii) *Output processes*, which determine how internal quantities are exposed to the rest of the network; (iii) *Connectivity functions*, which describe interactions with other populations within the same node, as well as external connections to other nodes and sensory inputs; and (iv) *Stochastic dynamics* of the model. This modular decomposition is particularly well-suited to neural mass models and provides a natural basis for formalizing model construction. To facilitate comparison, Table 1 presents the structural decomposition of the canonical models and identifies their major dynamical components.

Guided by this decomposition principle, we designed a probabilistic grammar to formalize the construction of node-level neural population models from these components. Table 2 summarizes the core production rules that define node-level model construction. The start symbol Node expands into intermediate nonterminal symbols (shown in blue), which are ultimately instantiated as terminal modelling choices (shown in orange). Expansion proceeds until only terminal symbols remain, yielding a complete structural specification of the node and a blueprint for assembling the corresponding dynamical system. Additionally, each production alternative is associated with a sampling probability. Rather than choosing these probabilities uniformly, we use empirical priors estimated from the relative frequencies of corresponding constructions observed across the set of canonical models. The full grammar, including all production rules, sampling probabilities, and mathematical definitions of each dynamical block, is provided in Supplementary Material C.

**Table 2.**
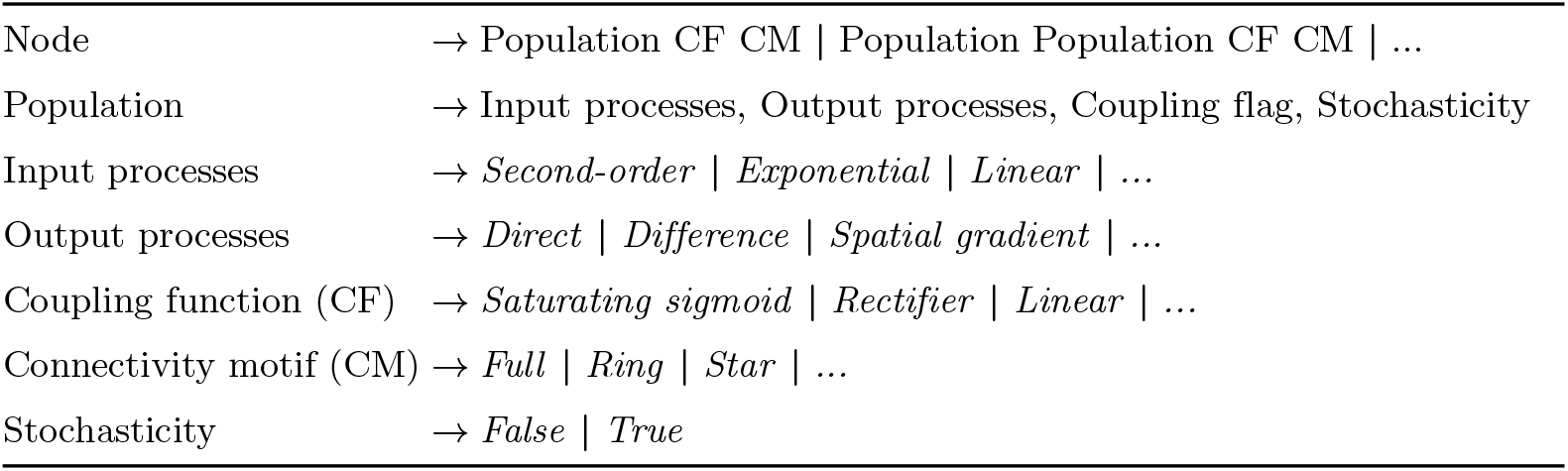
Simplified excerpt of the probabilistic grammar for node dynamics. It consists of nonterminal symbols, *terminal symbols* (shown in italics), and production rules describing how the starting nonterminal symbol Node is recursively expanded into terminal symbols corresponding to concrete model components. The grammar defines the hierarchical construction of nodes from populations, their input and output dynamics, intrinsic and extrinsic coupling functions (CF), connectivity motifs (CM), and stochastic noise. Vertical bars (|) indicate distinct alternatives within a rule. Only representative rules and structural alternatives are shown; the full grammar specification, including probabilities, is provided in Supplementary Material C.

To illustrate how a canonical neural mass model arises as a specific derivation within the grammar, we consider the classical Wilson–Cowan (WC) model. In this case, the Node symbol is expanded into two interacting populations. For each population, the grammar selects an exponential input process, a direct output process, and no stochasticity. The interaction between populations is specified by a baseline-subtracted sigmoidal coupling function, and the connectivity motif is instantiated as full, allowing reciprocal excitatory–inhibitory interactions. This yields a two-population architecture structurally equivalent to the standard WC model, up to parameter naming and notational conventions. A more detailed derivation of more complex canonical models, such as Jansen–Rit, is provided in Supplementary Material C.

More generally, a sampled grammar parse tree induces population-level dynamics of the following form. For each population *i* = 1, …, *n*_pops_ within a node, the grammar instantiates population-specific dynamics of the form

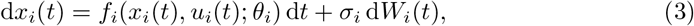

where *x*_*i*_(*t*) denotes the state vector of population *i, θ*_*i*_ its parameter set, and *W*_*i*_(*t*) a standard Wiener process with noise amplitude *σ*_*i*_ (with *σ*_*i*_ = 0 yielding a deterministic system). The functional form of *f*_*i*_ is determined by the grammar-selected input and output process blocks. The total input to population *i* is given by

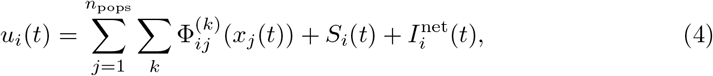

where 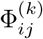 are grammar-selected coupling kernels, *S*_*i*_(*t*) denotes optional external or sensory input, and 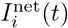 collects inputs from other nodes. Inter-regional interactions can include finite transmission delays by evaluating coupling terms on delayed states of other nodes, e.g., *x*_*j*_(*t* − *τ*_*ij*_). In the present work, however, 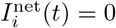, as we focus on single-node models. Equations (3)–(4) define the complete population-level dynamics instantiated by a grammar parse tree.

### Structural and empirical analysis of candidate models

Once generated, grammar-derived models were treated analogously to canonical models. Model complexity was quantified as the number of state equations, and structural proximity to canonical families was assessed using the composite structural distance metric already defined above. For a sampled model *M*, its distance to a canonical cluster *C*_*k*_ was defined as the minimum structural distance to any model in that cluster,

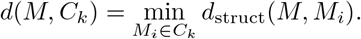

Each model was assigned to its closest cluster and characterized jointly by its complexity and minimal cluster distance. This representation places sampled models in a low-dimensional structural space defined by complexity and distance to the nearest canonical family, facilitating comparison, clustering, and identification of structurally novel or canonical-like models.

Empirical analysis followed the same general procedure as for canonical models: generated models were simulated, optimized, and evaluated using identical signal representations, loss functions, and evaluation metrics. The main difference concerned parameter initialization and optimization bounds. For canonical models, parameter defaults and search bounds were taken from model-specific values reported in the literature. For grammar-generated models, such model-specific prior information was not available. We therefore derived initial values and search bounds empirically, using descriptive statistics of fitted parameter values associated with each grammatical process across all canonical models and tasks in which that process appeared. To speed up these analyses, experiments were run in parallel on high-performance computing (HPC) clusters. The ENEEGMA source code is publicly available at https://github.com/NinaOmejc/ENEEGMA.

## Results

We first characterized the structural organization and empirical performance of canonical neural population models obtained from the literature, establishing a reference landscape in both structural space and empirical fit. We then evaluated grammar-derived candidate models within this landscape, assessing whether the probabilistic grammar generated structurally novel yet empirically competitive alternatives. Together, these analyses clarified how established models related to one another and whether the grammar meaningfully expanded the space of viable neural population dynamics.

### Structural analysis of canonical models

We first examined structural differences among canonical neural population models, quantifying variation in operator and function composition and symbolic structure. Structural difference was measured from the model equations, combining edit distance between symbolic expressions with cosine distance between operator-count vectors. The resulting 17 × 17 pairwise structural distance matrix is shown in Fig. 2A. Hierarchical clustering applied to the matrix produced a stable partition. Evaluation of mean silhouette scores across candidate cluster numbers (Fig. 2B) showed a maximum mean silhouette score of 0.48 at *k* = 6, supporting a six-cluster solution. The corresponding dendrogram in Fig. 2C delineates six structurally coherent model clusters.

**Fig 2.**
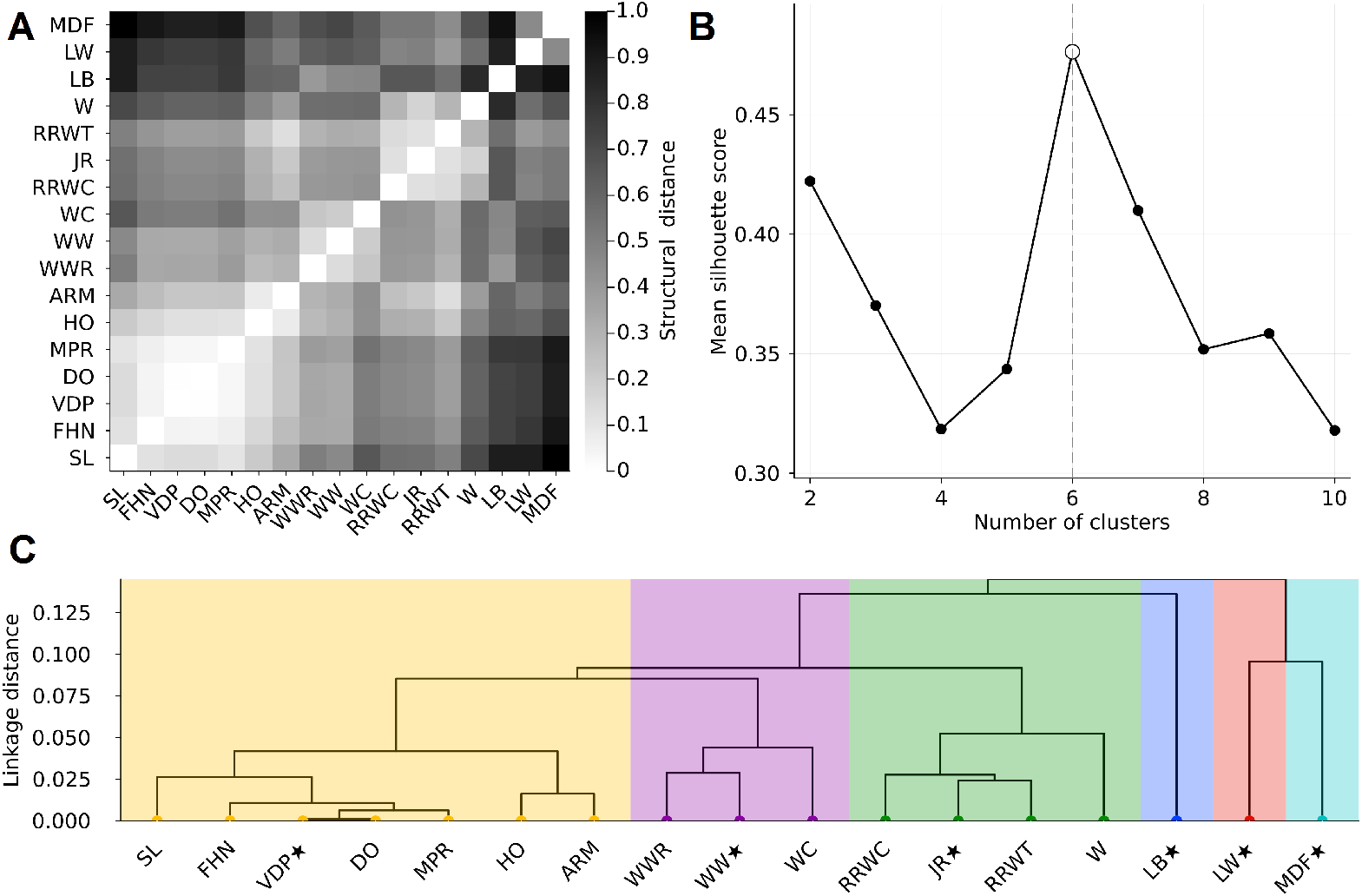
Structural similarity among canonical neural population models. (A) Pairwise structural distance matrix, where lower distances (lighter colors) indicate greater structural similarity between models. (B) Mean silhouette score as a function of the number of clusters, identifying an optimal partition at *k* = 6. (C) Hierarchical clustering based on structural distance separates the models into six structurally coherent clusters (colored regions). The black stars indicate cluster medoids, that is, the most centrally located models within each cluster, defined by the smallest average distance to all other models in the same cluster.

The largest cluster (color-coded in yellow) contains all phenomenological models and is dominated by low-order oscillator-like systems with a single population and polynomial nonlinearities, without explicit synaptic convolution or additional output processes. Notably, ARM also falls within this cluster despite having two populations and second-order dynamics. This likely reflects the fact that, in the present structural analysis, its relatively compact state space, linear coupling, and lack of sigmoidal transfer functions make it more similar to the low-order oscillator group than to more elaborate convolution-based neural mass models such as JR.

The second-largest cluster (color-coded in green) comprises higher-dimensional architectures organized around second-order synaptic kernels, including JR (medoid), W, and related RRW variants. Despite differences in connectivity motifs, these models share second-order filtered population dynamics and characteristic decay operators.

The third major cluster (color-coded in purple) is less homogeneous than the others and is defined more by a shared architectural template than by a single canonical mechanism. It comprises low-dimensional first-order population models with full coupling motifs and exponential-form transfer nonlinearities, including the baseline-subtracted sigmoid of WC and the relaxed rectifier used in WW/WWR. Lastly, MDF, LW, and LB form isolated branches. MDF is distinguished by its combination of multiple second-order synaptic subsystems and a difference-based output process, LW by a conductance-based membrane integrator layered on second-order kernels, and LB by voltage-dependent ionic currents and gating variables, placing it structurally apart from convolution-based neural mass formulations.

### Empirical analysis of canonical models

Having characterized the structural organization of canonical models, we next assessed their empirical fidelity to spectral features in EEG data.

Figure 3 and Table 3 summarize model performance, measured by the integrated absolute error (IAE) between empirical and simulated log-PSD, across four datasets per condition. For SSVEP, we report harmonic-restricted IAE (h-IAE) as the primary metric, since full-spectrum IAE can be dominated by broadband structure and thus understate stimulus-locked harmonic accuracy, which was our main quantity of interest. Because h-IAE is computed over a narrower frequency range, its absolute values are smaller and not directly comparable to RS IAE. We therefore base cross-condition comparison primarily on expected ranks.

**Table 3.** Empirical evaluation of canonical models via spectral fitting. Canonical models are ranked by their Bayesian expected rank *E*[rank], averaged across RS and SSVEP conditions. This rank-based summary is used because median integrated absolute error (IAE) values are condition-specific and therefore not directly comparable on a common scale across the two tasks. The table additionally reports model cluster, average, and condition-specific expected ranks, together with descriptive statistics: dataset-level median IAE for RS, harmonic-restricted IAE (h-IAE) for SSVEP, and relative interquartile range [Δ*Q*_1_, Δ*Q*_3_]. Lower values indicate better performance for both IAE and expected rank. Success rate denotes the percentage of runs that produced finite error values.

| Model | Cluster | E[rank] |  |  | RS IAE |  | SSVEP h-IAE |  | Success (%) |
| --- | --- | --- | --- | --- | --- | --- | --- | --- | --- |
| | | Avg. | RS | SSVEP | Med. | $[\Delta Q_1, \Delta Q_3]$ | Med. | $[\Delta Q_1, \Delta Q_3]$ | |
| MPR | 1 | 1.25 | 1.08 | 1.43 | 5.86 | [-0.08, 0.05] | 1.52 | [-0.55, 0.50] | 100.00 |
| FHN | 1 | 2.52 | 3.21 | 1.84 | 6.22 | [-0.06, 0.20] | 1.50 | [-0.18, 0.32] | 100.00 |
| SL | 1 | 3.13 | 3.06 | 3.20 | 6.27 | [-0.28, 0.33] | 1.87 | [-0.62, 0.61] | 100.00 |
| JR | 3 | 6.55 | 6.64 | 6.46 | 8.58 | [-1.54, 0.21] | 2.93 | [-0.41, 0.39] | 92.73 |
| WC | 2 | 6.96 | 10.14 | 3.77 | 8.29 | [-1.23, 1.70] | 1.86 | [-0.55, 0.94] | 90.77 |
| VDP | 1 | 8.30 | 3.47 | 13.13 | 6.10 | [-0.63, 0.70] | 2.88 | [-0.62, 3.00] | 100.00 |
| ARM | 1 | 8.32 | 4.68 | 11.97 | 6.56 | [-0.12, 0.32] | 3.85 | [-0.64, 0.80] | 100.00 |
| DO | 1 | 8.44 | 6.49 | 10.39 | 7.11 | [-0.34, 0.59] | 3.31 | [-0.43, 0.52] | 100.00 |
| RRWC | 3 | 8.83 | 10.71 | 6.95 | 9.78 | [-1.56, 1.37] | 2.99 | [-0.49, 0.43] | 53.33 |
| LB | 4 | 10.01 | 13.03 | 6.98 | 8.72 | [-0.68, 2.70] | 2.39 | [-0.72, 1.18] | 100.00 |
| LW | 5 | 10.21 | 12.45 | 7.98 | 10.7 | [-0.91, 0.70] | 2.96 | [-0.46, 0.47] | 33.33 |
| WWR | 2 | 11.04 | 14.41 | 7.66 | 11.5 | [-0.40, 0.28] | 3.18 | [-0.97, 0.78] | 100.00 |
| W | 3 | 11.05 | 9.59 | 12.51 | 8.28 | [-0.92, 0.95] | 3.98 | [-0.20, 0.29] | 87.69 |
| RRWT | 3 | 11.85 | 7.91 | 15.79 | 7.02 | [-0.29, 1.32] | 6.79 | [-1.10, 0.77] | 86.67 |
| WW | 2 | 12.31 | 13.13 | 11.49 | 11.5 | [-0.27, 2.40] | 3.82 | [-0.28, 0.38] | 100.00 |
| HO | 1 | 15.93 | 16.53 | 15.33 | 24.6 | [-1.46, 0.50] | 6.29 | [-0.63, 0.60] | 100.00 |
| MDF | 6 | 16.29 | 16.47 | 16.10 | 27.0 | [-3.70, 0.58] | 7.28 | [-1.30, 0.69] | 100.00 |

**Fig 3.**
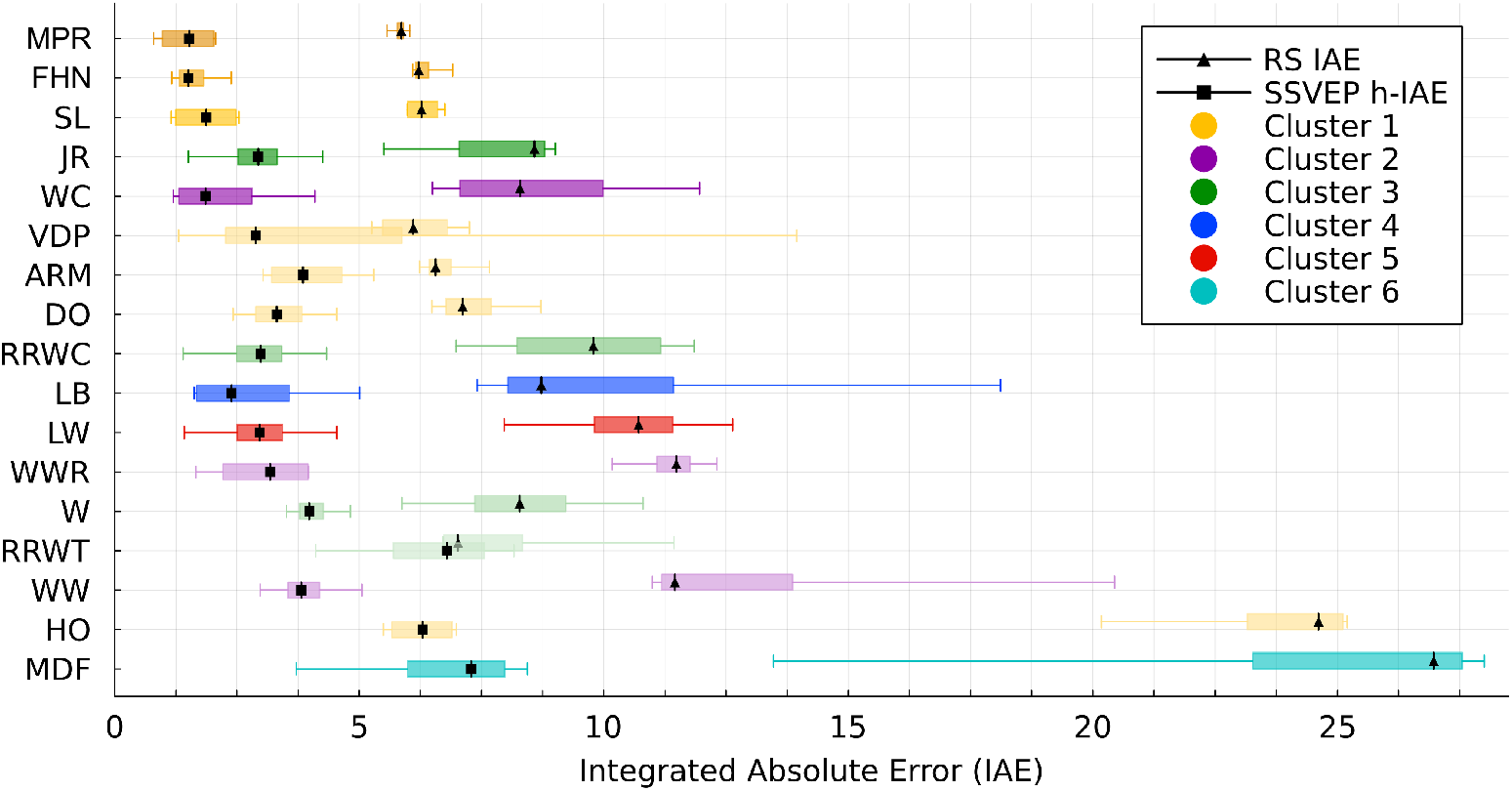
Canonical model comparison across RS and SSVEP conditions. Horizontal boxplots show IAE for RS data (with medians marked by triangles) and h-IAE for SSVEP data (with medians marked by squares). Models are ordered by their Bayesian expected rank, averaged across conditions.

Across both conditions, the MPR model achieved the strongest overall performance (*E*[rank] = 1.25) and the highest Bayesian posterior probability of being best (*P* (best) = 92.5% in RS and *P* (best) = 73.6% in SSVEP task). The second-best model, FHN, performed comparably well in the SSVEP condition (*E*[rank] = 1.84) but poorer for the RS. Together, these two models form the 95% cumulative credible set across both conditions, indicating that, given the observed data and bootstrap uncertainty, the posterior mass of the model ranking is concentrated on this pair. The third-ranked model, SL, with relatively high average rank (*E*[rank] = 3.13), also belongs to the phenomenological, low-order oscillator cluster. Together with MPR and FHN, it exhibits perfect numerical stability (100% success rate), reflecting robust performance across conditions. At a broader level, models in the phenomenological cluster were the strongest performers, particularly in the resting state.

Convolution-based synaptic-kernel architectures (e.g., JR, W, RRWT) occupied an intermediate performance range. Within this cluster, JR performed best, with an expected average rank of *E*[rank] = 6.55. Overall, these models reproduced oscillatory dynamics but exhibited substantially higher RS errors, suggesting a reduced ability to capture broadband spectral structure.

By contrast, several models ranked among the weakest performers.

Higher-dimensional neural population models such as LW and MDF showed elevated RS errors and received little posterior support (*E*[rank] *>* 12). LW was further limited by the lowest numerical stability of any model (33.33% success rate). HO, the structurally most constrained model, also performed poorly, with a high RS error (median IAE = 24.6) and an unfavorable expected rank (*E*[rank] = 15.93).

Figure 4 shows illustrative spectral fits for the three top-ranked canonical models together with the best-performing model from each structural cluster, thereby summarizing both the strongest overall performers and representative fits across model families. Results are shown only for the first dataset in each condition, which generally yielded the best fits, because hyperparameters were selected on this dataset and then fixed for the remaining ones. Fits for the other datasets and the remaining canonical models are provided in Supplementary Material D. Visual inspection of Fig. 4 indicates that the strongest phenomenological models, particularly MPR, FHN, and SL, achieved close resting-state fits, capturing both the alpha peak and the overall aperiodic decay with relatively low variability across repeated simulations; the largest residual discrepancies were concentrated mainly at low frequencies. SSVEP fits were generally more demanding, as accurate performance required matching several narrow stimulus-locked harmonic peaks. Even so, MPR, SL, and WC reproduced the harmonic structure well, whereas FHN and LB did so less consistently. By contrast, JR, MDF, and especially LW showed poorer and less stable SSVEP fits, with large run-to-run variability and, in some cases, non-finite error values.

**Fig 4.**
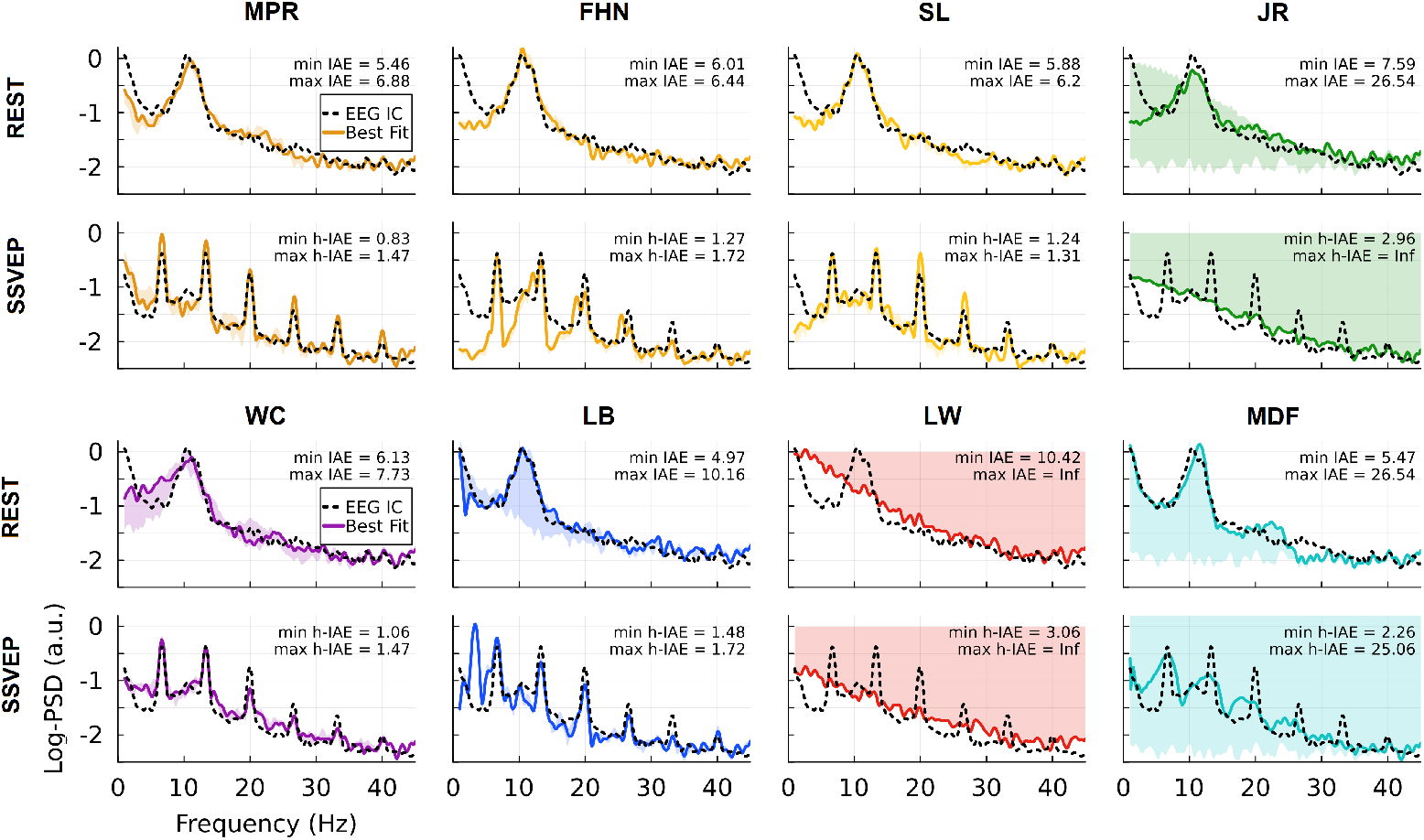
Log-PSD fits of selected canonical models for one representative dataset per condition. Panels show the three top-ranked canonical models together with the best-performing representative from each remaining structural cluster. Rows correspond to RS and SSVEP conditions. Black dashed lines denote empirical log-PSD, colored solid lines the best-fitting simulated spectra, and shaded regions the range (min–max) across five repeated simulations with different initial conditions and noise realizations. Insets report the minimum and maximum fitting error across repeated simulations (IAE for RS; h-IAE for SSVEP).

### Size and complexity of the grammar-generated model space

The grammar defines a rapidly expanding space of admissible node models (Table 4). Although the full grammar can generate a countably infinite number of models, it is still useful to obtain an approximate sense of the scale by treating the *polynomial* input dynamics option as a single admissible choice, rather than enumerating all of the infinite polynomial equations it could generate recursively. Under this simplification, the number of one-population variants follows directly from the available grammar choices for input processes, output processes, coupling terms, and stochasticity, yielding 432 within-population configurations. Combined with the admissible coupling functions and connectivity motif choices at the node level, this gives 5184 distinct one-population models. As the number of populations increases, the number of admissible models grows combinatorially through the combination of additional population blocks, coupling-function assignments, and larger connectivity-motif sets, reaching 1.96 × 10^17^ variants for *N* = 5.

**Table 4.** Combinatorial counts of node-model variants generated by the grammar as a function of the number of populations. Because the full grammar can generate a countably infinite number of models, the counts shown here are approximate and are obtained by treating the polynomial input dynamics option as a single admissible choice rather than recursively expanding all polynomial equations it could generate.

| Number of populations | Within-population configurations | Connectivity motifs | Coupling-function combinations | Total |
| --- | --- | --- | --- | --- |
| 1 | 432 | 2 | 6 | 5184 |
| 2 | 209,952 | 12 | 12 | $3.02 \times 10^7$ |
| 3 | $9.07 \times 10^7$ | 107 | 24 | $2.33 \times 10^{11}$ |
| 4 | $4.41 \times 10^{10}$ | 107 | 48 | $2.26 \times 10^{14}$ |
| 5 | $1.90 \times 10^{13}$ | 107 | 96 | $1.96 \times 10^{17}$ |

We additionally characterized one million grammar-generated models in terms of structural complexity. Figure 5 shows that the complexity of most sampled models lies within the range covered by the canonical models, as indicated by the dashed vertical lines, especially when complexity is quantified by the number of state equations. When quantified by the number of free parameters, the generated models still largely overlap the canonical range, but also extend more substantially beyond it.

**Fig 5.**
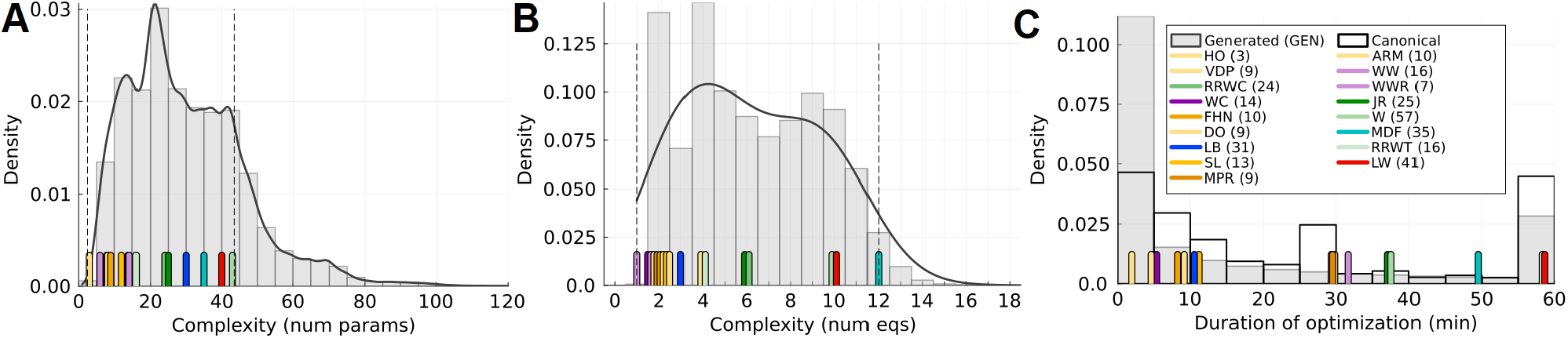
Structural complexity and optimization time for generated and canonical models. Histograms together with density curves show the complexity of one million models sampled from the probabilistic grammar, quantified either by the number of model parameters (Panel **A**) or by the number of state equations (Panel **B**). Color-coded ticks along the horizontal axis indicate the exact values for the canonical models listed in the legend, and dashed vertical lines mark the minimum and maximum values attained by the canonical set. A small horizontal jitter was added to the colored ticks in panel B for visual clarity. **C:** Optimization duration for a subset of 1000 generated models from the phenomenological, low-order oscillator-like region of the grammar-generated model space (GEN; gray histogram), compared with canonical models (white outlined histogram). In panel C, colored ticks indicate the median optimization duration of each canonical model, whereas the outlined histogram summarizes the full distribution across canonical models. Note that for practical computational reasons, the duration of each optimization run was limited to 60 minutes. Numbers in parentheses in the legend denote the number of parameters in each canonical model.

To assess computational cost, we separately examined optimization times for a subset of 1000 generated models drawn from the phenomenological, low-order oscillator-like region of the grammar-generated model space. Their generally short runtimes are consistent with the fact that the optimized generated subset was restricted to relatively simple, phenomenological low-order oscillator models. By contrast, the canonical models spanned a broader range of structural complexity, which was reflected in a wider distribution of optimization times, including substantially longer runs.

### Structural analysis of grammar-generated models

In Figure 6, we visualize the structural diversity of grammar-generated models. Each sampled model is assigned to one of the six canonical clusters based on its minimal structural distance to any model within that cluster. The models are embedded in a two-dimensional space defined by structural complexity (number of state equations) and structural distance to the nearest canonical model within the assigned cluster.

**Fig 6.**
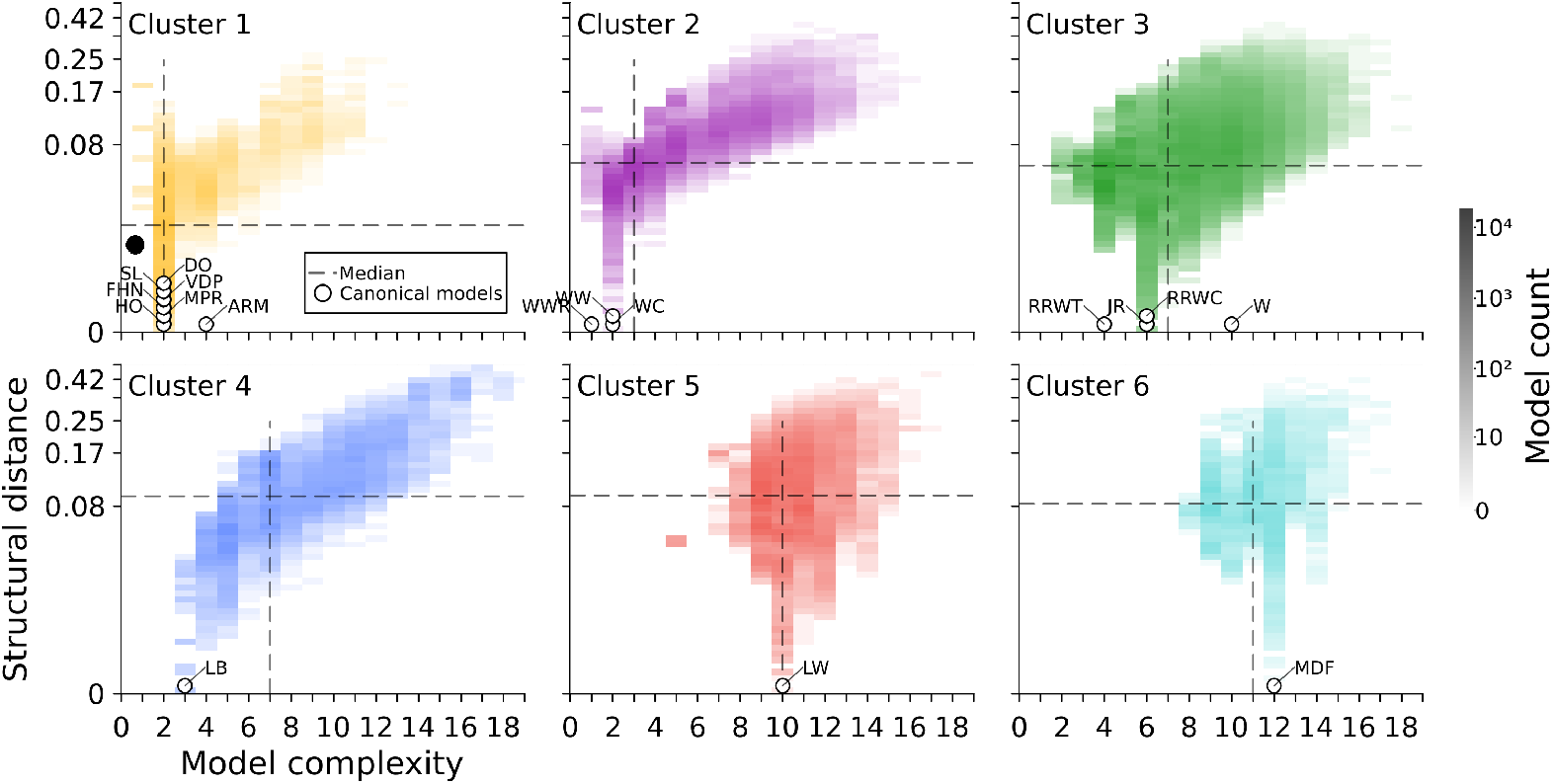
Grammar-sampled models in structural complexity–distance space. Each panel corresponds to one canonical model cluster. Generated models are assigned to clusters based on minimal structural distance to any canonical model within that cluster. The y-axis shows the structural distance to the nearest canonical model within the assigned cluster, and color intensity reflects model density. Dashed vertical and horizontal lines denote the median model complexity and median structural distance, respectively. White circles mark canonical models; their structural distance is zero and is shown with vertical jitter for visibility. The filled black circle indicates the sampled region for empirical evaluation, namely the lower-left quadrant of the best-performing Cluster 1 (yellow).

To aid interpretation, we divided the complexity–distance space into four quadrants using the median structural complexity and median structural distance within each panel (dashed lines in Fig. 6). For example, the upper-right quadrant contains highly complex and structurally distant models, which tend to be algebraically intricate and less tractable. In contrast, the lower-left quadrant contains simple models that remain close to canonical forms, making them a natural starting point for further evaluation because of their balance of interpretability and continuity with established formulations. Guided by this partition, we did not attempt to evaluate the full combinatorial space of grammar-generated models. Instead, we restricted sampling to the lower-left quadrant of the phenomenological, low-order oscillator-like cluster (color-coded in yellow), which showed the strongest empirical performance among the canonical model clusters. From this region, we empirically evaluated 1,000 models. This focused evaluation provides a preliminary exploration of the grammar-defined model space and serves as a proof of concept for the proposed framework.

### Functional analysis of grammar-generated models

Most of the evaluated 1,000 grammar-generated models produced numerically valid simulations, with success rates exceeding 75% in the RS condition and 95% in the SSVEP condition.

Table 5 summarizes the results at the cluster level. In Panel A, which compares the best-performing model within each cluster, the generated cluster, with its best model G1, ranked second overall (*E*[rank] = 1.67), behind canonical phenomenological Cluster 1, with the MPR model (*E*[rank] = 1.51). This difference was driven by resting-state performance, where canonical Cluster 1 performed best, whereas the generated cluster achieved the strongest SSVEP fits (*E*[rank] = 1.0).

**Table 5.** Cluster-level empirical performance of canonical and grammar-derived models. Panel A reports the best-performing model within each cluster, whereas Panel B reports the median performance across models within each cluster. Cluster groups are ranked according to their Bayesian expected rank (*E*[rank]), averaged across RS and SSVEP conditions, as well as across datasets. The table reports average and condition-specific Bayesian expected ranks, together with descriptive statistics: dataset-level median IAE (or h-IAE) and relative interquartile range [Δ*Q*_1_, Δ*Q*_3_]. Lower values indicate better performance for both error and rank. The top-performing cluster for each condition is bolded.

| Cluster | E[Rank] |  |  | RS IAE |  | SSVEP h-IAE |  |
| --- | --- | --- | --- | --- | --- | --- | --- |
| | Avg. | RS | SSVEP | Med. | $[\Delta Q_1, \Delta Q_3]$ | Med. | $[\Delta Q_1, \Delta Q_3]$ |
| *Panel A. Best model in cluster* |  |  |  |  |  |  |  |
| **Cluster 1 (MPR)** | **1.51** | **1.03** | 2.00 | **5.71** | **[-0.23, 0.20]** | 1.09 | [-0.12, 0.29] |
| **Generated (G1)** | 1.67 | 2.34 | **1.00** | 6.83 | [-0.34, 0.03] | **0.68** | **[-0.12, 0.11]** |
| Cluster 3 (JR) | 3.55 | 2.63 | 4.46 | 6.30 | [-0.52, 1.00] | 2.93 | [-0.44, 0.39] |
| Cluster 2 (WC) | 3.65 | 4.29 | 3.00 | 8.29 | [-1.23, 1.60] | 1.86 | [-0.55, 0.66] |
| Cluster 4 (LB) | 5.15 | 5.33 | 4.98 | 8.72 | [-0.68, 2.70] | 2.39 | [-0.72, 1.18] |
| Cluster 5 (LW) | 5.47 | 5.38 | 5.56 | 10.71 | [-0.91, 0.70] | 2.96 | [-0.46, 0.47] |
| Cluster 6 (MDF) | 7.00 | 7.00 | 7.00 | 26.97 | [-3.70, 0.58] | 7.29 | [-1.30, 0.69] |
| *Panel B. Cluster median* |  |  |  |  |  |  |  |
| **Generated** | **1.68** | 2.36 | **1.00** | 8.46 | [-0.89, 0.88] | **1.18** | **[-0.08, 0.14]** |
| **Cluster 1** | 1.75 | **1.00** | 2.50 | **6.44** | **[-0.19, 0.21]** | 2.82 | [-0.63, 0.38] |
| Cluster 4 | 4.19 | 4.85 | 3.54 | 8.72 | [-0.68, 2.70] | 2.39 | [-0.72, 1.18] |
| Cluster 3 | 4.28 | 2.66 | 5.90 | 8.31 | [-0.62, 0.89] | 3.34 | [-0.16, 0.33] |
| Cluster 5 | 4.32 | 4.82 | 3.81 | 10.71 | [-0.91, 0.70] | 2.96 | [-0.46, 0.47] |
| Cluster 2 | 4.78 | 5.30 | 4.26 | 11.12 | [-0.33, 0.23] | 3.15 | [-0.94, 0.77] |
| Cluster 6 | 7.00 | 7.00 | 7.00 | 26.97 | [-3.70, 0.58] | 7.29 | [-1.30, 0.69] |

Panel B shows cluster-median performance and therefore reflects typical performance across models within each cluster.Here, the generated cluster ranked first overall (*E*[rank] = 1.68), narrowly ahead of canonical Cluster 1 (*E*[rank] = 1.75). Again, the generated cluster performed best in SSVEP, while the canonical Cluster 1 retained the advantage in RS. Together, these results suggest that generated phenomenological-like models are particularly effective at reproducing stimulus-locked harmonic structure, whereas canonical Cluster 1 remains stronger for resting-state spectra.

Figures 7 and 8 together summarize the empirical performance of four top-performing generated models (G1, G2, G4, and G9). In comparison with canonical reference models (MPR, FHN, SL, and JR), the generated models generally showed higher RS error than the best canonical phenomenological models, although all outperformed JR. In SSVEP, however, all four generated models achieved lower median h-IAE than MPR and substantially outperformed the remaining canonical references, with comparatively tight distributions. The representative log-PSD fits clarify this pattern: in the RS condition, the generated models generally reproduced the dominant alpha peak and more faithfully captured the low-frequency part of the spectrum. However, the alpha peak was often less pronounced than in the empirical data and residual mismatches remained. In the SSVEP condition, by contrast, they reproduced the stimulus-locked harmonic peaks well, consistent with their low h-IAE values, while mismatches were more evident in broadband spectral regions between and beyond the harmonics. This pattern is expected because optimization was performed using the composite spectral loss, whereas final SSVEP comparison emphasized harmonic-restricted error (h-IAE), which specifically rewards fidelity to the evoked harmonic structure rather than to the full broadband spectrum. Together, these results show that compact grammar-generated models can produce physiologically plausible spectra and are particularly effective and robust in capturing stimulus-locked harmonic structure, while offering more limited gains for resting-state spectral fitting.

**Fig 7.**
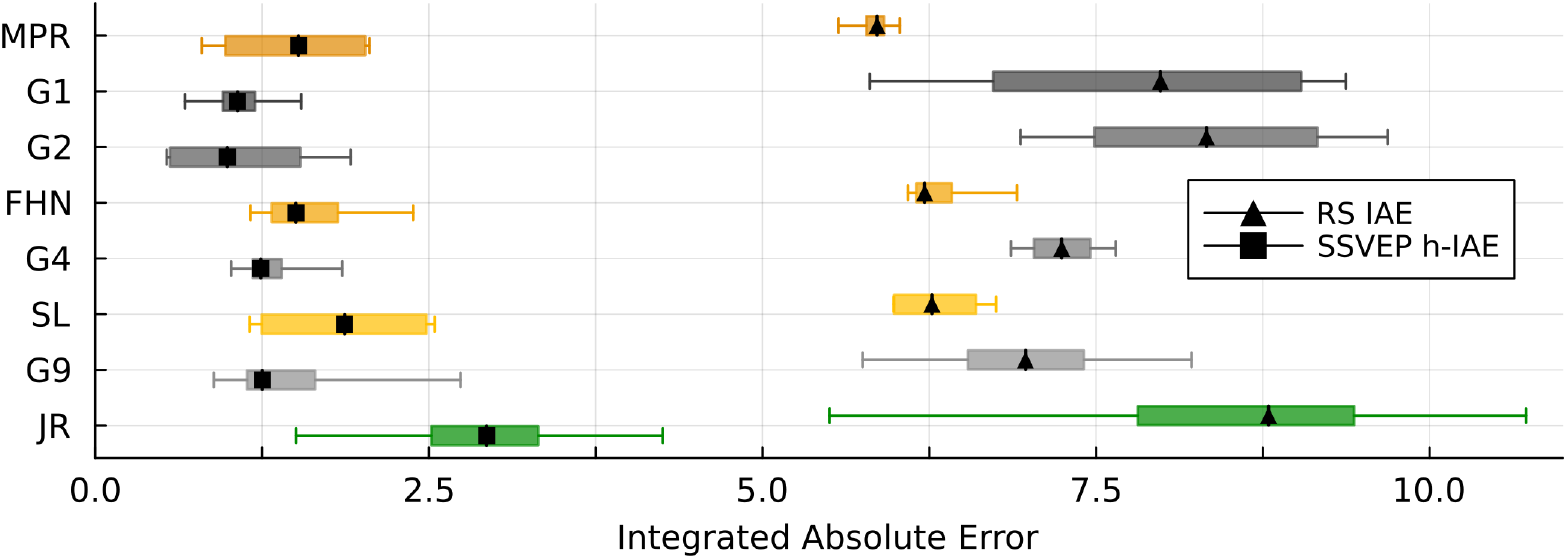
Empirical performance of four of the top-performing generated models. Horizontal boxplots show IAE for RS data (medians marked by triangles) and h-IAE for SSVEP data (medians marked by squares). Canonical models are included for reference. Models are ordered by their Bayesian expected rank, averaged across conditions. Although only a subset is shown here, rank computation included 20 generated models and 17 canonical models.

**Fig 8.**
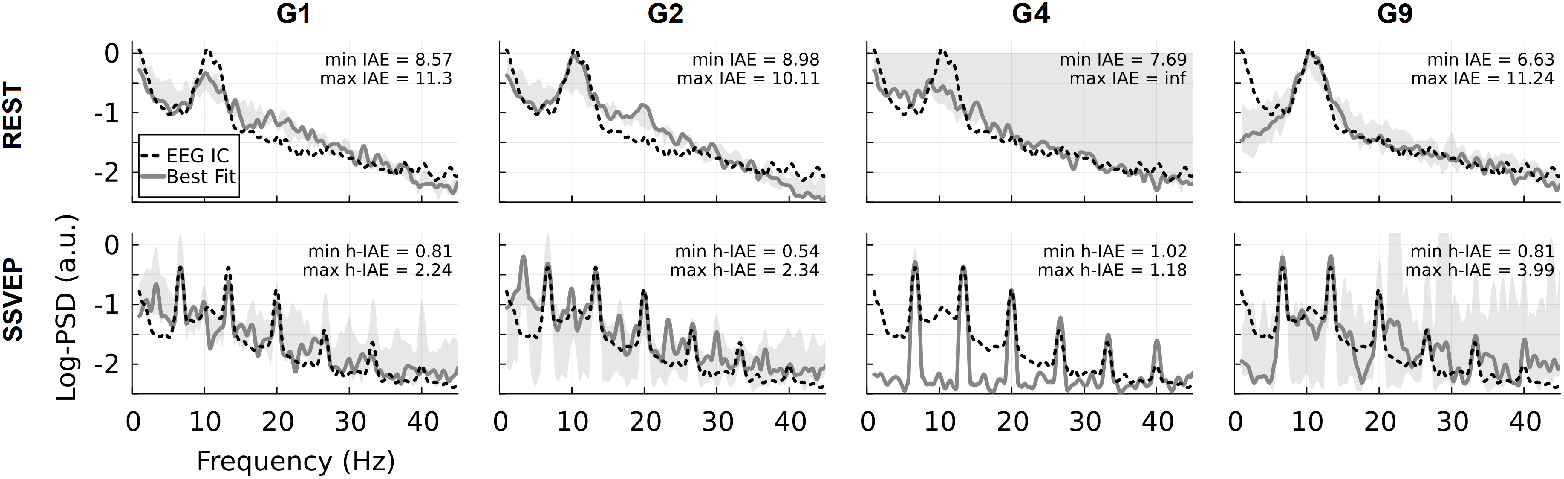
Log-PSD fits of best grammar-generated models. Columns show four representative top-performing generated models (G1, G2, G4, and G9), out of the 1,000 evaluated models in the GEN cluster, and rows correspond to RS and SSVEP conditions. The fits are shown for one representative dataset for each condition. Black dashed curves denote empirical log-PSD, solid gray curves the best-fitting simulated spectra, and shaded regions indicate the range (min–max) across five repeated simulations with varying initial conditions and noise realizations. Insets report the minimum and maximum fitting error across repeated simulations (IAE for RS; h-IAE for SSVEP). Because the SSVEP comparison was at the end based on harmonic-restricted error, close agreement is expected primarily at the stimulus-locked harmonic peaks rather than across the full broadband spectrum.

The selected generated models all belonged to a common class of compact two-dimensional polynomial, phenomenological systems with direct external drive and low-order nonlinear state interactions. As a representative example, model G1 is given by

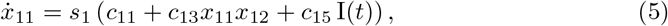

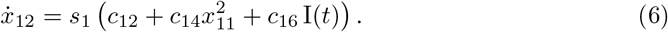

Here, *x*_11_(*t*) and *x*_12_(*t*) denote the two state variables, I(*t*) is the external input, *c*_*ij*_ are fitted coefficients controlling the constant, coupling, nonlinear, and input terms, and *s*_1_ is a global scaling parameter that sets the overall time scale of the dynamics. This system combines direct input to both state variables with bilinear coupling in 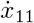 and quadratic self-feedback in 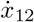, yielding a compact polynomial oscillator architecture that remains structurally simple while being flexible enough to reproduce the observed spectra. The remaining selected models, such as G4 and G9, can be understood as richer variants of the same general class, with additional higher-order and more asymmetric polynomial interactions. Complete equations and fitted parameter values are provided in Supplementary Material E.

## Discussion

In this study, we explored the space of neural population models with two main aims. First, we compared canonical models in terms of both structural organization and empirical performance on single-node EEG spectra. Second, we asked how distinctive these canonical formulations are within the broader model space, and whether systematic grammar-based exploration can reveal alternative node architectures with comparable or improved empirical fit.

To address these two questions, we first organized 17 canonical models within a common structural space derived from their symbolic equation structure and overall complexity. We then evaluated their empirical performance on EEG spectra from four datasets under two experimental conditions: resting state (RS) and steady-state visual evoked potentials (SSVEP). Furthermore, using canonical models as a basis, we constructed a probabilistic grammar, generated one million new candidate models, selected a subset of 1,000 models from a specific region of the structural space associated with the best-performing canonical model cluster, and assessed their performance on real EEG data. All analyses were performed within ENEEGMA, our custom Julia-based framework for grammar-based model construction and analysis. The main findings are discussed below.

### Among the canonical models, phenomenological, low-dimensional polynomial oscillators combine structural simplicity with the best single-node EEG spectral fits

The structural analysis showed that our 17 canonical models group into 6 clusters. In particular, 1) phenomenological, low-dimensional polynomial oscillators (VDP, SL, FHN, DO, MPR, ARM, HO) formed the largest cluster, distinct from 2) second-order synaptic-kernel neural mass models (JR, W, RRWT, RRWC), 3) low-dimensional population models with full coupling motifs and exponential-form transfer nonlinearities (WW, WWR, WC), and three additional single-model clusters: 4) LB, 5) LW, and 6) MDF.

Low-dimensional oscillator models, especially MPR, FHN, and SL, achieved the strongest overall performance across both RS and SSVEP conditions. This is not entirely surprising, given the long use of compact oscillator models and related low-dimensional dynamical systems in electrophysiological brain modelling (e.g. [30, 32, 76–78]), together with the fact that many salient EEG features are organized around rhythmic spectral components [1, 79]. Importantly, these models should not be understood as purely arbitrary phenomenological oscillators. MPR is a rigorous mean-field reduction of a spiking neuronal network [67], whereas FHN is a simplified two-dimensional model of neuronal excitability and spike generation, commonly viewed as a relaxation-oscillator reduction of Hodgkin–Huxley-type dynamics [65]. Their advantage in the present setting appears to lie in their compactness: they are sufficiently expressive to generate oscillatory peaks, waveform asymmetry, and harmonics, while remaining stable and relatively easy to optimize. By contrast, more mechanistically detailed neural mass models may encode richer biological structure, but that additional structure is not automatically rewarded in a single-node spectral fitting framework. Additionally, some models, such as WW and RWW, are typically used as local components of larger recurrent circuits or whole-brain systems [80], where oscillatory structure emerges through coupling, feedback, and network interactions rather than from isolated node dynamics alone [61]. Their relatively weak performance here is therefore not a general criticism of the Wong–Wang framework, but rather a consequence of the specific setting studied: single-node spectral fitting without inter-regional coupling. Likewise, the poorer performance of models such as LW, RRWC, and RRWT was not only a matter of fit quality but also of numerical robustness, indicating that stability under repeated optimization and simulation is itself an important practical dimension of model adequacy.

Taken together, these findings show that canonical neural population models differ in both structural organization and empirical performance in the single-node spectral setting. Their symbolic architecture is associated with systematic differences in empirical adequacy, with some model classes proving better matched to EEG spectra in the present setting than others.

### Grammar-based exploration can generate novel models with strong empirical performance

The ENEEGMA framework makes the node-model structure itself a search variable. Rather than selecting a fixed model family a priori and optimizing only its parameters, as in most current brain modeling toolboxes [36, 38, 39, 41, 81], the framework represents node models compositionally in terms of populations, input processes, output processes, coupling functions, connectivity motifs, and stochasticity. This provides a common and extensible language for comparing established neural population models and for generating new candidate systems from the same set of interpretable building blocks. Because the grammar is modular, it can also be adapted to specific scientific questions, for example, by adding new dynamical components, restricting particular constructions, or expanding the space of admissible mechanisms.

The empirical results from the grammar-derived models provide a proof of principle for this approach. Although the grammar defines a much larger admissible model space, the present study evaluated only 1,000 samples from one restricted region within the structural space of the best-performing phenomenological, low-dimensional cluster. Even under this limited sampling regime, the generated models were competitive with nearly all canonical families. At the cluster level, the best generated models ranked second overall and achieved the strongest SSVEP fits of all clusters. This is consistent with the preceding section, which showed that compact low-dimensional oscillator-like mechanisms are particularly effective in the present single-node spectral setting.

More broadly, the grammar is useful not only for generating new models, but also for asking which structural ingredients are actually needed to reproduce EEG spectra. Some of the best-performing generated systems remained structurally close to known oscillator forms, whereas others were less readily identifiable with canonical normal forms. This shows that grammar-based exploration can recover both canonical-like and structurally novel models with strong empirical performance.

### Spectral fit alone does not uniquely identify model structure

These findings also raise a broader methodological question: how distinguishable are alternative model structures when several of them can reproduce the same empirical spectra reasonably well? Previous work has largely examined parameter identifiability within fixed neural mass formulations [82–84]. Here, we highlight a complementary issue of model-level distinguishability: even when parameters are well fit, EEG spectra alone may remain insufficient to uniquely discriminate among structurally different population models. This limitation is evident in our results. Although model choice clearly mattered in the single-node setting studied here, several structurally different models still produced reasonably good fits, suggesting that viable EEG node models may be more numerous than is often assumed and that spectral adequacy alone is insufficient to identify a unique underlying mechanism or circuit architecture. The present findings, therefore, indicate that identification from EEG spectra alone may be intrinsically limited, particularly when optimization is based on high-level summary statistics such as the power spectral density. In that sense, a good spectral fit supports a plausible modelling description, but not necessarily a unique biological mechanism.

### Limitations and future work

Several limitations of the present study should be acknowledged. First, the empirical evaluation was based on four datasets and one representative independent component per condition, which limits both statistical power and the breadth of physiological variation. Future work should therefore extend the evaluation to larger cohorts, more components, and more diverse datasets. Nevertheless, the present study still represents a more substantial empirical test than is common in much of the neural population modelling literature, where models are often assessed in simulated settings and only rarely against realistic EEG recordings.

Second, only 1,000 generated models from one restricted region of the grammar-defined space were evaluated. The present search should therefore be regarded as preliminary rather than exhaustive. Moreover, because the best-performing canonical models belonged to the simplest phenomenological cluster, our targeted sampling focused on that part of the structural space rather than on more biologically detailed neural mass models, where the advantages of the probabilistic grammar may prove even greater. Future work will therefore require broader sampling across the admissible model space.

Third, all models were evaluated as isolated single nodes. This was a deliberate simplification, but, as already mentioned, it favors models that can generate relevant spectral structure locally. A natural next step is therefore to extend the framework to coupled multi-node and whole-brain settings, where the interaction between local dynamics and network coupling can be evaluated directly.

Fourth, the grammar currently does not include several important neural population models, including the Epileptor model [33, 85], next-generation neural mass models [6, 86], the Zerlaut mean-field model [87], low-dimensional density reductions by Pietras and Schwalger [88, 89], the Huang–Lin formulation [90], some extensions of the Jansen–Rit model [91], and phase-based dynamical models [92–94]. Some of these are not directly compatible with the current grammar because of fundamental differences in state representation, whereas others would require moderate or substantial extensions, for example, to support partial differential equations. Expanding the grammar to incorporate these families is therefore an important direction for future work.

Fifth, future work should employ more efficient search strategies, for example, using variational autoencoder-based generative models [51, 95]. These approaches use a trained neural generative model to concentrate sampling on more promising regions of model space, rather than relying on random sampling from the grammar.

Finally, we also plan to enable export of generated models in a TVB-compatible format. This would facilitate their use in whole-brain simulations and could improve parameter estimation by leveraging existing TVB optimisation tools [81, 96].

**In conclusion**, canonical neural population models were shown to be structurally organized and empirically distinguishable rather than interchangeable in the single-node spectral setting considered here. In particular, compact low-dimensional polynomial oscillator models such as MPR and FHN combined structural simplicity with the strongest overall EEG spectral fits. Second, grammar-based exploration showed that canonical formulations do not exhaust the space of viable EEG node models: even a restricted search recovered alternative architectures with competitive empirical performance. Together, these results establish an EEG-grounded framework for principled, comparative, and data-driven exploration of neural population models.

## Acknowledgments

We gratefully acknowledge Sebastian Mežnar for his help with context-dependent grammar sampling.

## Notes

https://github.com/NinaOmejc/ENEEGMA

### Competing Interest Statement

The authors have declared no competing interest.

### Funding

- The Slovenian Research and Innovation Agency: The research program “Knowledge Technologies” (P2-0103), The Gravity project “AI for Science” (GC-0001), The MAGMA project (N2-0378)

