## Supplementary Material for "Neural Population Models for EEG: From Canonical Models to Alternative Model Structures"

### Supplementary material A: Description of canonical models

Several of the neural mass models considered here originate from spatially extended mean-field or neural field formulations; in this work, we analyze their commonly used spatially lumped reductions.

This appendix summarizes the canonical neural-mass and phenomenological models used to construct the grammar. To provide a consistent representation, all models are rewritten in first-order form with state variables denoted $x_1, x_2, \ldots$. Model-specific parameters are retained in their original notation, while conceptually shared parameters (e.g. time constants, synaptic rates, coupling weights) are unified whenever this can be done without altering the meaning of the model.

**Wilson–Cowan model.**

$$\tau_e \dot{x}_1 = -x_1 + S(w_{ee}x_1 - w_{ei}x_2 + u_e(t)), \tag{1}$$
$$\tau_i \dot{x}_2 = -x_2 + S(w_{ie}x_1 - w_{ii}x_2 + u_i(t)), \tag{2}$$

with baseline subtracting sigmoid function:

$$S(v) = \frac{1}{1 + \exp[-a(v - \theta)]} - \frac{1}{1 + \exp(a\theta)}.$$

Here $x_1$ and $x_2$ denote excitatory and inhibitory firing rates, filtered by time constants $\tau_e$ and $\tau_i$. The four coupling parameters $w_{ee}, w_{ei}, w_{ie}, w_{ii}$ encode excitatory–excitatory, excitatory–inhibitory, inhibitory–excitatory, and inhibitory–inhibitory interactions, respectively, while $a$ and $\theta$ define the slope and threshold of the firing-rate nonlinearity. The terms $u_e(t)$ and $u_i(t)$ represent the total afferent input to the node.

**Alpha rhythm model.**

$$\mathrm{d}x_1 = x_2\,\mathrm{d}t, \tag{3}$$
$$\mathrm{d}x_2 = \big[A_e\alpha_e^2\big(u_e(t) - w_{ei}x_3\big) - 2\alpha_e x_2 - \alpha_e^2 x_1\big]\mathrm{d}t + \sigma_e\,\mathrm{d}W_{e,t}, \tag{4}$$
$$\mathrm{d}x_3 = x_4\,\mathrm{d}t, \tag{5}$$
$$\mathrm{d}x_4 = \big[A_i\alpha_i^2\big(u_i(t) + w_{ie}x_1\big) - 2\alpha_i x_4 - \alpha_i^2 x_3\big]\mathrm{d}t + \sigma_i\,\mathrm{d}W_{i,t}, \tag{6}$$

where $W_{e,t}$ and $W_{i,t}$ are independent Wiener processes. The variables $x_1$ and $x_3$ represent relay and reticular membrane potentials, and $x_2, x_4$ are their time derivatives introduced by rewriting the second-order alpha kernels into first-order form. Parameters $\alpha_e, \alpha_i$ determine synaptic rates, $A_e, A_i$ scale PSP amplitudes, and $w_{ie}, w_{ei}$ are excitatory–inhibitory loop weights. The inputs $u_e(t)$ and $u_i(t)$ denote pooled afferent drives to the relay and reticular populations, while $\sigma_e$ and $\sigma_i$ control the amplitude of stochastic perturbations that sustain narrowband alpha oscillations.

**Jansen–Rit model.**

$$\dot{x}_1 = x_2, \tag{7}$$
$$\dot{x}_2 = Aa\,S(x_3 - x_5) - 2a\,x_2 - a^2 x_1, \tag{8}$$
$$\dot{x}_3 = x_4, \tag{9}$$
$$\dot{x}_4 = Aa\big(u(t) + C_2 S(w_e x_1)\big) - 2a\,x_4 - a^2 x_3, \tag{10}$$
$$\dot{x}_5 = x_6, \tag{11}$$
$$\dot{x}_6 = Bb\,S(w_i x_1) - 2b\,x_6 - b^2 x_5, \tag{12}$$

with

$$S(v) = \frac{2e_0}{1 + \exp[r(v_0 - v)]}.$$

State variables $(x_1, x_2)$ correspond to the pyramidal (PYR) population postsynaptic potential and its derivative; $(x_3, x_4)$ correspond to the excitatory interneuron (EIN); and $(x_5, x_6)$ correspond to the inhibitory interneuron (IIN). Parameters $A$ and $B$ set excitatory and inhibitory synaptic gains, while $a$ and $b$ define their characteristic decay rates. The coupling weights $w_e$ and $w_i$ replace the original $C_1$ and $C_3$ and encode the strength of PYR→EIN and PYR→IIN projections, respectively. The remaining gain $C_2$ modulates the EIN→PYR feedback. The external drive $u(t)$ represents the pooled afferent input to the column, such as thalamic excitation or the exogenous sensory input.

**Wendling–Chauvel model.**

$$\begin{aligned}
\dot{x}_1 &= x_2, \\
\dot{x}_2 &= Aa\, S(x_3 - x_5 - x_7) - 2ax_2 - a^2 x_1, \\
\dot{x}_3 &= x_4, \\
\dot{x}_4 &= Aa\big(p(t) + C_1 S(w_e x_1)\big) - 2ax_4 - a^2 x_3, \\
\dot{x}_5 &= x_6, \\
\dot{x}_6 &= Bb\, S(w_s x_1) - 2bx_6 - b^2 x_5, \\
\dot{x}_7 &= x_8, \\
\dot{x}_8 &= Gg\, S(w_f x_1) - 2gx_8 - g^2 x_7, \\
\dot{x}_9 &= x_{10}, \\
\dot{x}_{10} &= Hh\, S(w_a x_7) - 2hx_{10} - h^2 x_9,
\end{aligned}$$

with

$$S(v) = \frac{2e_0}{1 + \exp[r(v_0 - v)]}.$$

In this form, $(x_1, x_2)$ represent the pyramidal population (PYR) postsynaptic potential and its derivative; $(x_3, x_4)$ represent the excitatory interneuron (EIN); $(x_5, x_6)$ the slow dendritic inhibitory interneuron; $(x_7, x_8)$ the fast somatic inhibitory interneuron; and $(x_9, x_{10})$ an additional inhibitory interneuron mediating slow–fast feedback. Parameters $A, B, G, H$ and rates $a, b, g, h$ define the amplitudes and time constants of the four synaptic kernels. The coupling weights $w_e, w_s, w_f$ describe, respectively, the strength of excitatory, slow inhibitory, and fast inhibitory projections to the pyramidal population, while $w_a$ encodes the coupling from the fast inhibitory population to the auxiliary interneuron. The remaining constant $C_1$ encodes intrinsic feedback pathways within the interneuron network.

**Moran–David–Friston model.**

The equations below follow the reduced first-order formulation summarized and implemented by [1], rather than a direct transcription of the original DCM papers.

$$
\begin{aligned}
\dot{x}_1 &= x_2, && (13)\\
\dot{x}_2 &= H_e S(x_7) - 2a_e x_2 - a_e^2 x_1 + \eta(t), && (14)\\
\dot{x}_3 &= x_4, && (15)\\
\dot{x}_4 &= H_e S(x_1) - 2a_e x_4 - a_e^2 x_3, && (16)\\
\dot{x}_5 &= x_6, && (17)\\
\dot{x}_6 &= H_i S(x_9) - 2a_i x_6 - a_i^2 x_5, && (18)\\
\dot{x}_7 &= x_8, && (19)\\
\dot{x}_8 &= H_e S(x_7) - 2a_e x_8 - a_e^2 x_7, && (20)\\
\dot{x}_9 &= x_{10}, && (21)\\
\dot{x}_{10} &= H_i S(x_9) - 2a_i x_{10} - a_i^2 x_9, && (22)\\
\dot{x}_{11} &= x_8 - x_{10}, && (23)
\end{aligned}
$$

with

$$S(v) = \frac{1}{1+\exp[-\kappa(v-\theta)]} - \frac{1}{1+\exp(\kappa\theta)}.$$

In this reordered formulation, $(x_1, x_2)$, $(x_3, x_4)$, $(x_5, x_6)$, $(x_7, x_8)$, and $(x_9, x_{10})$ correspond to the membrane potentials and their derivatives of the excitatory and inhibitory subpopulations across superficial and deep cortical layers. The parameters $H_e$ and $H_i$ are excitatory and inhibitory synaptic gains, while $a_e$ and $a_i$ determine the associated time constants. The observable $x_{11}$ yields a layer-weighted pyramidal output. The sigmoid parameters $\kappa$ and $\theta$ determine the slope and threshold of the population firing-rate nonlinearity.

**Liley–Wright model.**

$$
\begin{aligned}
\dot{x}_1 &= x_2, && (24)\\
\dot{x}_2 &= -2\gamma_{ee} x_2 - \gamma_{ee}^2 x_1 + \gamma_{ee}\Gamma_{ee}\, A\big(x_5; N_{ee}^{\mathrm{b}}, p_{ee}, S_{e,\max}, \mu_e, \sigma_e\big), && (25)\\
\dot{x}_3 &= x_4, && (26)\\
\dot{x}_4 &= -2\gamma_{ei} x_4 - \gamma_{ei}^2 x_3 + \gamma_{ei}\Gamma_{ei}\, A\big(x_5; N_{ei}^{\mathrm{b}}, p_{ei}, S_{e,\max}, \mu_e, \sigma_e\big), && (27)\\
\tau_e \dot{x}_5 &= h_{e,r} - x_5 + \Psi(x_5; h_{ee,\mathrm{eq}}, h_{e,r})\, x_1 + \Psi(x_5; h_{ie,\mathrm{eq}}, h_{e,r})\, x_3, && (28)\\
\dot{x}_6 &= x_7, && (29)\\
\dot{x}_7 &= -2\gamma_{ie} x_7 - \gamma_{ie}^2 x_6 + \gamma_{ie}\Gamma_{ie}\, A\big(x_{10}; N_{ie}^{\mathrm{b}}, p_{ie}, S_{i,\max}, \mu_i, \sigma_i\big), && (30)\\
\dot{x}_8 &= x_9, && (31)\\
\dot{x}_9 &= -2\gamma_{ii} x_9 - \gamma_{ii}^2 x_8 + \gamma_{ii}\Gamma_{ii}\, A\big(x_{10}; N_{ii}^{\mathrm{b}}, p_{ii}, S_{i,\max}, \mu_i, \sigma_i\big), && (32)\\
\tau_i \dot{x}_{10} &= h_{i,r} - x_{10} + \Psi(x_{10}; h_{ei,\mathrm{eq}}, h_{i,r})\, x_6 + \Psi(x_{10}; h_{ii,\mathrm{eq}}, h_{i,r})\, x_8, && (33)\\
& && (34)
\end{aligned}
$$

All reversal-potential scalings share a unified form

$$\Psi(v; h_{\mathrm{eq}}, h_r) = \frac{h_{\mathrm{eq}} - v}{|h_{\mathrm{eq}} - h_r|},$$

and the effective presynaptic rates are written as

$$A(v; N^{\mathrm{b}}, p, S_{\max}, \mu, \sigma) = N^{\mathrm{b}}\, S(v; S_{\max}, \mu, \sigma) + p,$$

with a shared sigmoidal firing-rate function

$$S(v; S_{\max}, \mu, \sigma) = \frac{S_{\max}}{1 + \exp\big(-\sqrt{2}\,(v-\mu)/\sigma\big)}.$$

We represent the excitatory population by states $(x_1, \ldots, x_5)$ and the inhibitory population by $(x_6, \ldots, x_{10})$, with the membrane potentials placed last in each population. Here $x_5$ and $x_{10}$ are the excitatory and inhibitory membrane potentials. The variables $(x_1, x_2)$, $(x_3, x_4)$, $(x_6, x_7)$ and $(x_8, x_9)$ form second-order synaptic kernels for the *ee*, *ei*, *ie*, and *ii* pathways. Parameters $\tau_e, \tau_i$ are membrane time constants, $h_{e,r}, h_{i,r}$ resting potentials, and the pairs $(\Gamma_{..}, \gamma_{..})$ set synaptic strength and decay. The terms $N^{\mathrm{b}}_{..}$ and $p_{..}$ are background and constant presynaptic rates. This reduced Liley–Wright formulation preserves the conductance-based structure of the original model while remaining low-dimensional.

**Robinson–Rennie–Wright thalamocortical model.**

For the cortical node we use

$$\dot{x}_{c1} = x_{c2}, \tag{35}$$

$$\dot{x}_{c2} = -(\alpha+\beta)x_{c2} + \alpha\beta\big(-x_{c1} + \nu_{ee}x_{c3} + \nu_{ei}S(x_{c1}) + \nu_{es}S(x_{t1}(t-\tau))\big), \tag{36}$$

$$\dot{x}_{c3} = x_{c4}, \tag{37}$$

$$\dot{x}_{c4} = -\gamma x_{c4} - \gamma^2 x_{c3} + \gamma^2 S(x_{c1}), \tag{38}$$

$$\dot{x}_{c5} = x_{c6}, \tag{39}$$

$$\dot{x}_{c6} = -(\alpha+\beta)x_{c6} + \alpha\beta\big(-x_{c5} + \nu_{re}x_{c3}(t-\tau) + \nu_{rs}S(x_{c5})\big), \tag{40}$$

with

$$S(v) = \frac{Q_{\max}}{1 + \exp[-(v-\theta)/\sigma]}.$$

Here $x_{c1}$ denotes the cortical excitatory membrane potential and $x_{c3}$ the cortical excitatory axonal field, with $x_{c2}$ and $x_{c4}$ their first derivatives introduced by the first-order reduction of the original second-order equations. The pair $(x_{c5}, x_{c6})$ represents a cortical inhibitory population and its derivative. The parameters $\alpha, \beta$ set the synaptic rise and decay rates, $\gamma$ governs axonal propagation, and the gains $\nu_{ee}, \nu_{ei}, \nu_{es}$ specify local excitatory coupling and delayed thalamocortical input.

For the thalamic node we use

$$\dot{x}_{t1} = x_{t2}, \tag{41}$$

$$\dot{x}_{t2} = -(\alpha+\beta)x_{t2} + \alpha\beta\big(-x_{t1} + \nu_{se}x_{c3}(t-\tau) + \nu_{sr}S(x_{t3}) + \nu_{sn}\Phi_n(t)\big), \tag{42}$$

$$\dot{x}_{t3} = x_{t4}, \tag{43}$$

$$\dot{x}_{t4} = -(\alpha+\beta)x_{t4} + \alpha\beta\big(-x_{t3} + \nu_{re}x_{c3}(t-\tau) + \nu_{rs}S(x_{t1})\big), \tag{44}$$

with the same sigmoid $S$.

In the thalamic subsystem, $x_{t1}$ and $x_{t3}$ represent the specific and reticular thalamic membrane potentials, while $x_{t2}$ and $x_{t4}$ are their derivatives. The gains $\nu_{se}$ and $\nu_{re}$ encode delayed corticothalamic input from the cortical excitatory field $x_{c3}$, $\nu_{sr}$ and $\nu_{rs}$ describe intra-thalamic coupling, and $\nu_{sn}\Phi_n(t)$ represents a subcortical (noise) drive. In ENEEGMA we treat these cortical and thalamic subsystems as two separate nodes that together form the full RRW thalamocortical unit.

**Wong–Wang excitatory–inhibitory model.**

These are TVB implementations of the reduced Wong–Wang family, rather than direct transcriptions of the original 2006 decision model [2].

$$\mathrm{d}x_1 = \left( -\frac{x_1}{\tau_1} + (1 - x_1)\,\gamma_1\,S(w_p J_N x_1 - J_i x_2 + u_1(t)) \right) \mathrm{d}t + \sigma_1\,\mathrm{d}W_{1,t}, \tag{45}$$

$$\mathrm{d}x_2 = \left( -\frac{x_2}{\tau_2} + \gamma_2\,S(J_N x_1 - x_2 + u_2(t)) \right) \mathrm{d}t + \sigma_2\,\mathrm{d}W_{2,t}. \tag{46}$$

with relaxed-rectifier transfer

$$S(I) = \frac{aI - b}{1 - \exp[-d(aI - b)]}.$$

Here $x_1$ and $x_2$ denote the excitatory and inhibitory synaptic gating fractions. The parameters $\tau_1$ and $\tau_2$ are synaptic decay time constants, while $\gamma_1$ and $\gamma_2$ denote population gain factors. The parameters $w_p$, $J_N$, and $J_i$ encode recurrent excitatory and inhibitory coupling, and $u_1(t)$ and $u_2(t)$ represent constant external drive to the two populations.

The function $S(\cdot)$ is a parametric approximation of the firing-rate input–output relationship derived from noisy integrate-and-fire neurons in Wong and Wang (2006). The parameters $a$, $b$, and $d$ control the shape of the nonlinearity. The noise amplitudes $\sigma_1$ and $\sigma_2$ scale independent Wiener processes $W_{1,t}$ and $W_{2,t}$; in the deterministic limit, the model is recovered by setting $\sigma_1 = \sigma_2 = 0$.

**Reduced Wong–Wang model.** This is a one-dimensional reduced Wong–Wang model, from the TVB implementation [3].

$$\mathrm{d}x_1 = \left( -\frac{x_1}{\tau_s} + (1 - x_1)\,\gamma\,S\big(wJ_N x_1 + u(t)\big) \right) \mathrm{d}t + \sigma\,\mathrm{d}W_t, \tag{47}$$

with a nonlinear transfer

$$S(I) = \frac{aI - b}{1 - \exp[-d(aI - b)]}.$$

Here, $x_1$ is the excitatory synaptic gating fraction. The term $wJ_N x_1$ represents recurrent excitation, and $u(t)$ denotes pooled external input (which in classical formulations reduces to a constant baseline $I_0$). The parameters $\tau_s$ and $\gamma$ are the synaptic decay constant and gain, respectively. The constants $a, b, d$ determine the shape of the relaxed-rectifier nonlinearity, and $\sigma$ sets the amplitude of additive stochastic fluctuations via the Wiener process $W_t$.

**Larter–Breakspear model.**

$$\dot{x}_1 = -\Big( I_{\mathrm{Ca}}(x_1) + I_{\mathrm{K}}(x_1, x_2) + I_{\mathrm{L}}(x_1) + I_{\mathrm{Na}}(x_1) + I_{\mathrm{IE}}(x_1, x_3) \Big) + a_{ne}\,u(t), \tag{48}$$

$$\dot{x}_2 = \phi\,\frac{m_{\mathrm{K}}(x_1) - x_2}{\tau_{\mathrm{K}}}, \tag{49}$$

$$\dot{x}_3 = b\big(a_{ni}\,u(t) + a_{ei}\,x_1\,Q_V(x_1)\big), \tag{50}$$

where the ionic and synaptic currents are

$$I_{\mathrm{Ca}}(x_1) = \big(g_{\mathrm{Ca}} + (1-C)\, r_{\mathrm{NMDA}}\, a_{ee}\, Q_V(x_1)\big)\, m_{\mathrm{Ca}}(x_1)\, \big(x_1 - V_{\mathrm{Ca}}\big), \tag{51}$$

$$I_{\mathrm{K}}(x_1, x_2) = g_{\mathrm{K}}\, x_2\, \big(x_1 - V_{\mathrm{K}}\big), \tag{52}$$

$$I_{\mathrm{L}}(x_1) = g_{\mathrm{L}}\, \big(x_1 - V_{\mathrm{L}}\big), \tag{53}$$

$$I_{\mathrm{Na}}(x_1) = \big(g_{\mathrm{Na}}\, m_{\mathrm{Na}}(x_1) + (1-C)\, a_{ee}\, Q_V(x_1)\big)\, \big(x_1 - V_{\mathrm{Na}}\big), \tag{54}$$

$$I_{\mathrm{IE}}(x_1, x_3) = a_{ie}\, x_3\, Q_Z(x_3). \tag{55}$$

The voltage-dependent activation curves and population firing rates are

$$m_{\mathrm{ion}}(x) = \tfrac{1}{2}\Big(1 + \tanh \frac{x - T_{\mathrm{ion}}}{d_{\mathrm{ion}}}\Big), \qquad \mathrm{ion} \in \{\mathrm{Ca}, \mathrm{Na}, \mathrm{K}\}, \tag{56}$$

$$Q_V(x_1) = Q_{V,\max}\, \tfrac{1}{2}\Big(1 + \tanh \frac{x_1 - V_T}{d_V}\Big), \tag{57}$$

$$Q_Z(x_3) = Q_{Z,\max}\, \tfrac{1}{2}\Big(1 + \tanh \frac{x_3 - Z_T}{d_Z}\Big). \tag{58}$$

Here $x_1$ is the pyramidal membrane potential, $x_2$ the potassium activation variable, and $x_3$ an inhibitory interneuron drive. The parameters $g_{\mathrm{Ca}}, g_{\mathrm{K}}, g_{\mathrm{L}}, g_{\mathrm{Na}}$ are maximal calcium, potassium, leak, and sodium conductances with corresponding reversal potentials $V_{\mathrm{Ca}}, V_{\mathrm{K}}, V_{\mathrm{L}}, V_{\mathrm{Na}}$. The thresholds $T_{\mathrm{Ca}}, T_{\mathrm{Na}}, T_{\mathrm{K}}$ and widths $d_{\mathrm{Ca}}, d_{\mathrm{Na}}, d_{\mathrm{K}}$ shape the ionic activation curves, while $V_T, Z_T$ and $d_V, d_Z$ play the same role for the population firing rates $Q_V$ and $Q_Z$, scaled by $Q_{V,\max}$ and $Q_{Z,\max}$. Synaptic strengths $a_{ee}, a_{ei}, a_{ie}, a_{ne}, a_{ni}$ control excitatory and inhibitory coupling, $r_{\mathrm{NMDA}}$ weights the NMDA contribution, and $C$ sets the AMPA–NMDA mixing. The constants $\tau_{\mathrm{K}}$ and $\phi$ govern potassium kinetics, and $b$ sets the time scale of the inhibitory drive $x_3$. The term $u(t)$ denotes the external input (corresponding to $I_{\mathrm{ext}}$ in the original formulation). This conductance-based mean-field model combines ionic and synaptic nonlinearities and can generate bursting, fast oscillations, and chaotic dynamics.

**Harmonic oscillator model.**

$$\dot{x}_1 = x_2, \tag{59}$$

$$\dot{x}_2 = -2\zeta\omega_0 x_2 - \omega_0^2 x_1 + u(t). \tag{60}$$

The variable $x_1$ represents displacement or membrane potential, with $x_2$ its time derivative. The parameters $\omega_0$ and $\zeta$ are the natural frequency and damping ratio; $u(t)$ is an external drive.

**Montbrió–Pazó–Roxin model.** MPR is special — it is a rigorous mean-field reduction, but functionally it behaves like a phenomenological oscillator at the population level.

$$\dot{x}_1 = \frac{\Delta}{\pi} + 2x_1 x_2, \tag{61}$$

$$\dot{x}_2 = x_2^2 + \eta_0 - (\pi x_1)^2 + u(t), \tag{62}$$

where $x_1$ denotes population firing rate and $x_2$ the mean membrane potential of a QIF network. The parameters $\Delta$ and $\eta_0$ set intrinsic heterogeneity and mean drive.

**FitzHugh–Nagumo model.**

$$\dot{x}_1 = x_1 - \frac{1}{3}x_1^3 - x_2 + u(t), \tag{63}$$

$$\dot{x}_2 = \frac{1}{\tau}(a + x_1 - bx_2). \tag{64}$$

The activator variable $x_1$ interacts with a slow recovery variable $x_2$ through parameters $a, b$, with $\tau$ governing the time-scale separation.

**Van der Pol model.**

$$\dot{x}_1 = x_2, \tag{65}$$

$$\dot{x}_2 = \mu(1 - x_1^2)x_2 - x_1. \tag{66}$$

Here $x_1$ is the oscillator position and $x_2$ its velocity; $\mu$ determines the strength of nonlinear damping.

**Stuart–Landau model.**

$$\dot{x}_1 = \mu x_1 - \omega x_2 - (x_1^2 + x_2^2)x_1, \tag{67}$$

$$\dot{x}_2 = \omega x_1 + \mu x_2 - (x_1^2 + x_2^2)x_2. \tag{68}$$

This system is the real two-dimensional form of the Hopf normal form. The parameter $\mu$ controls the stability and amplitude of oscillations, while $\omega$ sets the intrinsic frequency. For $\mu > 0$, trajectories converge to a limit cycle of radius $\sqrt{\mu}$, and the dynamics reduce asymptotically to a phase oscillator. In this sense, the Stuart–Landau model is a state-space extension of simple phase-dynamics models, augmenting phase evolution with an explicit amplitude degree of freedom.

**Duffing Oscillator.**

$$\dot{x}_1 = x_2, \tag{69}$$

$$\dot{x}_2 = -\alpha x_1 - \beta x_1^3 - 2\delta x_2 + F(t). \tag{70}$$

The system models a nonlinear oscillator with linear stiffness $\alpha$, cubic stiffness $\beta$, damping $\delta$, and forcing $F(t)$. It is a standard benchmark for nonlinear and chaotic dynamics.

### Supplementary material B: Details on data preprocessing

Empirical EEG data were obtained from the dataset of Lee et al. [4], accessed via the MOABB framework [5]. The 64-channel recordings were converted to MATLAB format (R2023a, The MathWorks, USA) and preprocessed in EEGLAB [6]. Continuous data were resampled to 256 Hz, 60 Hz line noise was attenuated, and the signals were band-pass filtered between 0.1 and 48 Hz. Noisy channels were detected automatically using the Clean Rawdata plug-in (version 2.0) with a flatline criterion of 10 s, a channel-correlation criterion of 0.85, and a maximum tolerated bad-time fraction of 0.5, and were then interpolated using spherical interpolation. The data were additionally inspected manually, and bad segments were removed. Recordings were rereferenced to the common average, and independent component analysis was performed using extended Infomax (`runica`). Component source localization was further characterized using DIPFIT with a standard BEM head model, MNI coordinates, and the standard 10–05 electrode layout. Components were classified with ICLabel, and non-brain or artifactual components were rejected semi-automatically using a probability threshold of 0.3 for eye components and 0.5 for the remaining non-brain classes, followed by additional manual inspection. On average, we retained X brain components. For the SSVEP condition, stimulus events were relabeled according to the four stimulation frequencies (5.45, 6.67, 8.57, and 12.0 Hz), and the data were epoched from −500 to 4500 ms relative to stimulus onset, with baseline correction over −200 to 0 ms. To obtain representative neural signals for model fitting, components classified as brain were clustered across datasets using spectral, scalp-topographic, dipole-location, and dipole-moment features. When multiple components from the same dataset fell into the same cluster, the component with the lowest dipole residual variance was retained. From these clustered components, one representative resting-state component and one representative SSVEP component at 6.67 Hz were selected for each of the four analyzed datasets.

### Supplementary material C: Details on probabilistic grammar

We first present the full grammar in Table 1, then describe its components, and finally illustrate the model generation process with a detailed example of the Jansen-Rit model.

The resulting grammar defines a structured yet flexible generative model of node dynamics, balancing expressiveness with fidelity to established neural-mass formulations. Each terminal symbol in the grammar corresponds to a dynamical building block. Chosen blocks are assembled automatically into population-level and node-level dynamical systems. All equations below use the standard notation $x(t)$ for state variables, $u(t)$ for the pooled presynaptic drive, and $\theta$ for parameters.

#### Input processes

Input process blocks transform a pooled presynaptic firing-rate signal $u(t)$ into low-dimensional internal state variables. In the grammar, each block is an ODE system of the form

$$\dot{\mathbf{x}}(t) = f(\mathbf{x}(t), u(t); \theta), \tag{71}$$

and populations may have one or two such blocks, depending on the grammar parse tree. We consider five explicit input-process families: linear, exponential, gating-kinetics, second-order PSP, and voltage-dependent dynamics, together with one grammar-generated polynomial family. These six families were selected because, across canonical neural mass and phenomenological models (Table **??**), every population-level synaptic or intrinsic subsystem can be represented or closely approximated by one of these dynamical templates. Grammar-generated polynomial dynamics is the most flexible and provides a broad basis for data-driven discovery. The remaining families provide a compact yet mechanistically interpretable basis that spans classical EEG modelling practice. Below, we list mathematical descriptions of the input process blocks:

**Linear dynamics.** This block represents the simplest affine input-driven dynamics, with baseline term $a$ and input gain $b$:

$$\dot{x}(t) = a + b\,u(t). \tag{72}$$

**Table 1.** Probabilistic grammar for node dynamics. It is composed of nonterminal symbols, *terminal symbols*, and production rules, which describe the evolution of the starting nonterminal symbol Node to the terminal symbols, which represent concrete model choice. Vertical bars (|) indicate distinct alternatives within a rule. Each production alternative is associated with a probability (shown in square brackets), enabling stochastic sampling of node architectures. The grammar defines the hierarchical construction of nodes from populations (Pop1, Pop2), their intrinsic and extrinsic coupling functions (ECF, PCF, CF), input and output processes (InputProcess, OutputProcess), algebraic combinations of variables (S, P, V), and connectivity motifs (CM, Digit). Abbreviations: Pop = population; ECF = extrinsic coupling function; PCF = population coupling function; CF = connectivity function; CM = connectivity motif; S = sum; P = product; V = variable.

| | | | | | | |
|---|---|---|---|---|---|---|
| Node | → | Pop1 CF SCF CM1 | | | | [0.50] |
| | | Pops2 CF SCF{1..2} CM2 | | | | [0.28] |
| | | Pop1 Pops2 CF SCF{1..3} CM | | | | [0.06] |
| | | Pops2 Pops2 CF SCF{1..4} CM | | | | [0.06] |
| | | Pops2 Pops2 Pop1 CF SCF{1..5} CM | | | | [0.10] |
| Pops2 | → | Pop1 Pop1 | [0.78] | \| | Pop2 | [0.22] |
| Pop1 | → | InputProcess OutputProcess ECF ECF Stochasticity | | | | [1.00] |
| Pop2 | → | InputProcess InputProcess OutputProcess ECF ECF ECF ECF Stochasticity | | | | [1.00] |
| InputProcess | → | S S | [0.14] | \| | *Second-order* | [0.66] |
| | | *Exponential* | [0.08] | \| | *Gating kinetics* | [0.06] |
| | | *Linear* | [0.03] | \| | *Voltage-dependent* | [0.03] |
| OutputProcess | → | *Direct readout* | [0.84] | \| | *Membrane integrator* | [0.06] |
| | | *Difference* | [0.06] | \| | *Spatial gradient* | [0.04] |
| SCF | → | *Custom* | [0.89] | \| | *Linear* | [0.11] |
| ECF | → | *Linear* | [0.53] | \| | *False* | [0.32] |
| | | *Custom* | [0.15] | | | |
| CF | → | *Saturating sigmoid* | [0.52] | \| | *Baseline-subtracted sigmoid* | [0.24] |
| | | *Relaxed rectifier* | [0.24] | | | |
| S | → | P + S | [0.5] | \| | P | [0.5] |
| P | → | V * P | [0.42] | \| | V | [0.58] |
| V | → | *x1* | [0.56] | \| | *x2* | [0.44] |
| CM1 | → | *Null* | [0.8] | \| | *Full* | [0.2] |
| CM2 | → | *Full* | [0.50] | \| | *Ring* | [0.30] |
| | | Digit | [0.20] | | | |
| CM | → | *Full* | [0.06] | \| | *Ring* | [0.06] |
| | | *Star* | [0.40] | \| | *Hub-tail* | [0.06] |
| | | *Star-tail* | [0.06] | \| | *Star-loop-extended* | [0.15] |
| | | *EI-extended* | [0.15] | \| | *Erdos–Renyi* Digit Digit | [0.06] |
| Digit | → | *0* | [0.1] | \| | *1* | [0.1] |
| | | *2* | [0.1] | \| | *3* | [0.1] |
| | | *4* | [0.1] | \| | *5* | [0.1] |
| | | *6* | [0.1] | \| | *7* | [0.1] |
| | | *8* | [0.1] | \| | *9* | [0.1] |
| Stochasticity | → | *False* | [0.83] | \| | *True* | [0.17] |

**First-order exponential synaptic kernel:**

$$\dot{x}(t) = \frac{1}{\tau}\left(u(t) - \kappa\, x(t)\right) \tag{73}$$

The parameters are the effective time constant $\tau > 0$ and the decay factor $\kappa > 0$. It is also called a linear leaky integrator and is the canonical Wilson–Cowan synaptic filter.

**First-order gating kinetics.**

$$\dot{x}(t) = -\frac{1}{\tau}x(t) + \kappa\,(1 - x(t))\, u(t) \tag{74}$$

It represents a synaptic gating variable that opens proportionally to the input $u(t)$, and relaxes back with time constant $\tau > 0$ and input gain $\kappa > 0$. Used in Wong–Wang synaptic gating models [2, 3, 7].

**Second-order PSP kernel.**

$$\dot{x}_1 = x_2, \tag{75}$$
$$\dot{x}_2 = \omega^2\big(g\,u(t) - x_1\big) - 2\zeta\omega\,x_2, \tag{76}$$

where $\omega > 0$ is the natural frequency, $g$ an input gain, and $\zeta$ is the damping ratio. All second-order PSP variants (bi-exponential, $\alpha$-kernel, and resonant kernels) are represented within this single damped-driven oscillator family. Special cases include when $\zeta = 1$, we get a critically damped $\alpha$-kernel, and when $\zeta > 1$, we get over-damped bi-exponential kernels. When $\zeta < 1$, we get resonant kernels (harmonic oscillator). For reference, the classical *alpha kernel* with amplitude $A$ and rate $a > 0$ is

$$\dot{x}_1(t) = x_2(t), \tag{77}$$
$$\dot{x}_2(t) = Aa\,u(t) - 2a\,x_2(t) - a^2 x_1(t), \tag{78}$$

whose impulse response is proportional to $te^{-at}$. This is recovered from the unified form by identifying $g = A$, $\omega = a$, and $\zeta = 1$.

A more general *bi-exponential* kernel, with separate rise and decay time constants $\alpha, \beta > 0$, obeys

$$\dot{x}_1(t) = x_2(t), \tag{79}$$
$$\dot{x}_2(t) = \frac{1}{\alpha\beta}\big(u(t) - x_1(t)\big) - \Big(\frac{1}{\alpha} + \frac{1}{\beta}\Big)x_2(t), \tag{80}$$

which corresponds to a particular choice of $(\omega, \zeta, g)$ in the unified second-order system (here $g = 1$, and $\omega, \zeta$ are chosen so that $\omega^2 = 1/(\alpha\beta)$ and $2\zeta\omega = 1/\alpha + 1/\beta$). This unified second-order form covers the kernel dynamics used in JR, ARM, RRW, W, and LW, and also includes the harmonic oscillator as a special mathematical case.

**Voltage-dependent conductance-based dynamics.** The conductance-based block consists of three state variables: membrane potential $x_1(t)$, potassium gating variable $x_2(t)$, and an auxiliary inhibitory-state variable $x_3(t)$. The dynamics are

$$\dot{x}_1(t) = -\big(I_{\mathrm{Ca}}(t) + I_{\mathrm{K}}(t) + I_{\mathrm{L}}(t) + I_{\mathrm{Na}}(t) + I_{\mathrm{IE}}(t)\big) + I_{\mathrm{ext}}(t), \tag{81}$$

$$\dot{x}_2(t) = \phi\,\frac{m_{\mathrm{K}}(x_1(t)) - x_2(t)}{\tau_{\mathrm{K}}}, \tag{82}$$

$$\dot{x}_3(t) = b\left(a_{ni}\,u(t) + a_{ei}\,x_1(t)\,Q_V(x_1(t))\right), \tag{83}$$

where the ionic and synaptic currents are

$$I_{\mathrm{Ca}}(t) = \Big(g_{\mathrm{Ca}} + (1 - C)\,r_{\mathrm{NMDA}}\,a_{ee}\,Q_V(x_1(t))\Big)\,m_{\mathrm{Ca}}(x_1(t))\,(x_1(t) - V_{\mathrm{Ca}}), \tag{84}$$
$$I_{\mathrm{K}}(t) = g_{\mathrm{K}}\,x_2(t)\,(x_1(t) - V_{\mathrm{K}}), \tag{85}$$
$$I_{\mathrm{L}}(t) = g_{\mathrm{L}}\,(x_1(t) - V_{\mathrm{L}}), \tag{86}$$
$$I_{\mathrm{Na}}(t) = \Big(g_{\mathrm{Na}}\,m_{\mathrm{Na}}(x_1(t)) + (1 - C)\,a_{ee}\,Q_V(x_1(t))\Big)\,(x_1(t) - V_{\mathrm{Na}}), \tag{87}$$
$$I_{\mathrm{IE}}(t) = a_{ie}\,x_3(t)\,Q_Z(x_3(t)), \tag{88}$$
$$I_{\mathrm{ext}}(t) = a_{ne}\,u(t). \tag{89}$$

The voltage-dependent activation functions are

$$m_{\mathrm{ion}}(x) = \frac{1}{2}\left(1 + \tanh\frac{x - T_{\mathrm{ion}}}{d_{\mathrm{ion}}}\right), \qquad \mathrm{ion} \in \{\mathrm{Ca}, \mathrm{Na}, \mathrm{K}\}, \tag{90}$$

and the firing-rate transforms are

$$Q_V(x_1) = Q_{V,\max}\frac{1}{2}\left(1+\tanh\frac{x_1-V_T}{d_V}\right), \qquad Q_Z(x_3) = Q_{Z,\max}\frac{1}{2}\left(1+\tanh\frac{x_3-Z_T}{d_Z}\right). \tag{91}$$

Parameters $g_{\mathrm{Ca}}, g_{\mathrm{K}}, g_{\mathrm{L}}, g_{\mathrm{Na}}$ are the maximal calcium, potassium, leak, and sodium conductances, and $V_{\mathrm{Ca}}, V_{\mathrm{K}}, V_{\mathrm{L}}, V_{\mathrm{Na}}$ are the corresponding reversal potentials. The activation thresholds $T_{\mathrm{Ca}}, T_{\mathrm{K}}, T_{\mathrm{Na}}$ and widths $d_{\mathrm{Ca}}, d_{\mathrm{K}}, d_{\mathrm{Na}}$ determine the sigmoidal activation functions $m_{\mathrm{Ca}}, m_{\mathrm{K}}, m_{\mathrm{Na}}$. The firing-rate transforms $Q_V$ and $Q_Z$ are controlled by their maximal amplitudes $Q_{V,\max}$ and $Q_{Z,\max}$, thresholds $V_T$ and $Z_T$, and widths $d_V$ and $d_Z$. Synaptic coupling parameters $a_{ee}$, $a_{ei}$, $a_{ie}$, $a_{ne}$, and $a_{ni}$ govern, respectively, excitatory-to-excitatory, excitatory-to-inhibitory, inhibitory-to-excitatory, external-to-excitatory, and external-to-inhibitory influences. The parameter $r_{\mathrm{NMDA}}$ scales the NMDA-mediated component of excitatory input, while $C$ controls the AMPA–NMDA mixing ratio. Finally, $\tau_{\mathrm{K}}$ is the potassium gating time constant, $\phi$ sets the rate of potassium activation, and $b$ determines the time scale of the inhibitory-state variable $x_3$.

This three-variable conductance-based system corresponds to the dynamics of the Larter–Breakspear model. We do not treat the inhibitory-state variable $x_3$ as a separate population because, in the Larter–Breakspear formulation, it does not represent an autonomous neuronal subpopulation with its own incoming and outgoing synaptic projections. Instead, it acts as an internal slow modulatory variable contributing to inhibitory feedback through the nonlinear transform $Q_Z(x_3)$. Treating $x_1$, $x_2$, and $x_3$ as components of a single conductance-based population, therefore preserves the original mean-field interpretation and is consistent with common usage of the model in TVB and related literature.

**Grammar-generated polynomials.** Polynomial input processes are generated by the following recursive subgrammar:

$$\begin{aligned}
\texttt{InputProcess} \to &\quad S\,S \quad [0.14] \\
S \to &\quad S+P\,[0.50] \quad | \quad P\,[0.50] \\
P \to &\quad V*P\;[0.42] \quad | \quad V\,[0.58] \\
V \to &\quad x_1 \quad\;\; [0.56] \quad | \quad x_2[0.44]
\end{aligned}$$

It can construct polynomial expressions on the right-hand side of neural population equations. It produces exactly two expressions, since `InputProcess` expands to two instances of $S$. The nonterminal $S$ generates sums of multiplicative terms ($P$), and $P$ produces products of state variables ($V$). Rule probabilities are given in square brackets and were inferred empirically for terminal symbols.

Notably, the grammar-generated polynomial family is strictly more expressive than the other types. It can reproduce the structure of low-order polynomial oscillators and approximate some specialized dynamics, including second-order kernels. This overlap does not negate the value of keeping explicit kernel families in the grammar, as they encode the canonical PSP structures used throughout neural mass modelling and thereby provide strong mechanistic inductive biases and a constrained, interpretable search space. On the other hand, polynomial dynamics offer a flexible universal expansion that complements rather than replaces these physiologically grounded components. When counting the number of admissible models, we exclude the recursive polynomial subgrammar, which in principle generates a countably infinite family of polynomial input processes.

#### Output processes

Output processes determine how internal variables of a population are turned into presynaptic signals that are transmitted to other populations or nodes. In ENEEGMA, these are implemented as optional low-dimensional ODE blocks that append additional state variables to the population and mark a subset of them as "output" variables. For a given population, $\mathbf{x}(t)$ collects the variables created by the input process blocks, and $\mathbf{y}(t)$ contains the additional state variables introduced by an output process block. For notational brevity, when a single internal component is referenced without specifying an index, we write it as $x(t)$; otherwise, $x_j(t)$ denotes the $j$th component of the internal state vector $\mathbf{x}(t)$. When an output process introduces auxiliary state variables, these obey a system of differential equations of the generic form

$$\dot{\mathbf{y}}(t) = g(\mathbf{y}(t), \mathbf{x}(t); \boldsymbol{\phi}) \,, \tag{92}$$

where $\boldsymbol{\phi}$ are the parameters. In our grammar, we set four output process blocks:

**Direct readout.**

The simplest case, *direct readout*, introduces no additional state variables; instead, an existing internal variable is designated as the population's presynaptic output. This behaviour matches the WC, JR, W, MDF, WW, HO, and polynomial model families, which do not employ additional output processes.

**Spatial-gradient (second-order) dynamics.**

$$\dot{y}_1(t) = y_2(t), \tag{93}$$

$$\dot{y}_2(t) = \gamma^2(-y_1(t) + x(t)) - 2\gamma\, y_2(t), \tag{94}$$

where $\gamma > 0$ is a rate parameter and $x(t)$ is a designated internal state variable from the population's input process blocks. This subsystem implements a critically damped second-order filter acting on $x(t)$, providing temporal smoothing and shaping of the outgoing activity. Such construction appears in the RRW cortical model.

**Membrane integrator.**

$$\tau\, \dot{y}(t) = h_r - y(t) + \sum_j \Psi(y(t), h_r)\; x_j(t), \tag{95}$$

with the reversal-potential transform

$$\Psi(y(t), h_r) = \frac{h_r - y(t)}{|h_r - h_{\mathrm{eq}}|}. \tag{96}$$

Here $\tau > 0$ is a membrane time constant, $h_r$ a resting potential, and $x_j(t)$ are the input signals from the population's internal dynamics. The factor $\Psi$ scales each input according to the difference between the current membrane state $y(t)$ and the reversal potential $h_{\mathrm{eq}}$. This block captures a simplified conductance-based membrane integration, and is used for conductance-driven populations (e.g., the Larter–Breakspear model).

**Difference readout.**

$$\dot{y}(t) = x_i(t) - x_j(t), \tag{97}$$

where $x_i(t)$ and $x_j(t)$ are two designated internal state variables of the two input process blocks within the same population. The resulting state $y(t)$ accumulates the difference between the two designated internal variables and is exported as the population output. This construction is used in the MDF model.

#### Connectivity functions

Connectivity functions transform inputs into the actual drive that enters a population's state equations. Although functions such as sigmoids or rectifiers are sometimes described as "output transformations" in the neural-mass literature—because they map membrane potentials to firing rates—here we reserve the term *output processes* exclusively for mechanisms that introduce additional state variables. Connectivity functions, in contrast, apply transformations directly to incoming signals already appearing in the population's differential equations. Each population may assign different connectivity functions to inputs coming from other populations, sensory channels, or internode connections. For a given population, all incoming signals are grouped by the connectivity function $c$ assigned to them. The total transformed input is

$$h(t) = \sum_{c \in \mathcal{C}} h_c\big(x_c(t); \phi_c\big) + I(t), \tag{98}$$

where $\mathcal{C}$ is the set of all connectivity-function types and $\phi_c$ are the corresponding parameters. For each connectivity-function group $c$, the aggregated drive is

$$x_c(t) = \sum_{s \in \mathcal{U}_c} v_s\, u_s(t) + \sum_{s \in \mathcal{Z}_c} w_s\, z_s(t), \tag{99}$$

where $\mathcal{U}_c$ denotes the set of internode inputs processed by connectivity function $c$, and $\mathcal{Z}_c$ denotes the set of within-node population outputs processed by the same function. Here, $u_s(t)$ are internode inputs, $z_s(t)$ are outputs of other populations within the node, and $v_s$ and $w_s$ are the corresponding coupling weights. If present, the global sensory input $I(t)$ is added separately to the appropriate population input. Currently implemented connectivity functions are:

**Linear.**

$$h(x) = x. \tag{100}$$

A direct linear pass-through of the aggregated input, without parameters.

**Saturating logistic sigmoid.**

$$h(x) = \frac{e}{1 + \exp\big(\rho(\theta - x)\big)}. \tag{101}$$

A classical logistic nonlinearity with maximum value $e$, slope parameter $\rho$, and midpoint (threshold) $\theta$. Implements the firing-rate transfer used in many neural-mass models.

**Baseline-subtracted logistic sigmoid.**

$$h(x) = g\left(\frac{1}{1 + \exp\big(a(\theta - x)\big)} - \frac{1}{1 + \exp(a\theta)}\right). \tag{102}$$

A logistic sigmoid shifted so that $h(0) = 0$. Parameter $a$ controls steepness, $\theta$ the midpoint, and $g$ is an overall gain.

**Relaxed rectifier (soft threshold).**

$$h(x) = \frac{ax - b}{1 - \exp\big(-d(ax - b)\big)}. \tag{103}$$

A smooth approximation of a rectified linear response. Parameter $a$ is an input gain, $b$ a threshold, and $d$ controls the softness of the transition.

#### Connectivity motifs

Connectivity motifs specify the adjacency pattern among populations within a node. A full binary connectivity matrix on $n$ populations has $n^2$ possible directed edges, so the number of distinct adjacency patterns grows as $2^{n^2}$. For $n = 5$, this already yields $2^{25} \approx 3.4 \times 10^7$ possibilities, which would make connectivity choices dominate the grammar's model space. To avoid this combinatorial explosion and to bias models toward interpretable structures, we restrict the grammar to a finite library of reusable connectivity motifs.

For $n \leq 2$, the grammar can enumerate all sensible $n \times n$ patterns (i.e. those that actually connect the populations). For $n > 2$, instead of sampling an arbitrary binary matrix, the grammar selects one of several predefined motifs and instantiates it at size $n$. These motifs are implemented as Boolean adjacency masks and fall into two broad categories: *deterministic* motifs, which come from canonical models and graph theory, and the seeded *random* motifs, which generate adjacency patterns from standard network models, e.g., Erdos–Renyi networks.

Motifs drastically reduce the model space compared to sampling arbitrary adjacency matrices: for an $n$-population node, a free adjacency matrix has $2^{n^2}$ possibilities, while a motif chooses one of $\mathcal{O}(10)$ families with fixed or parameterised structure.

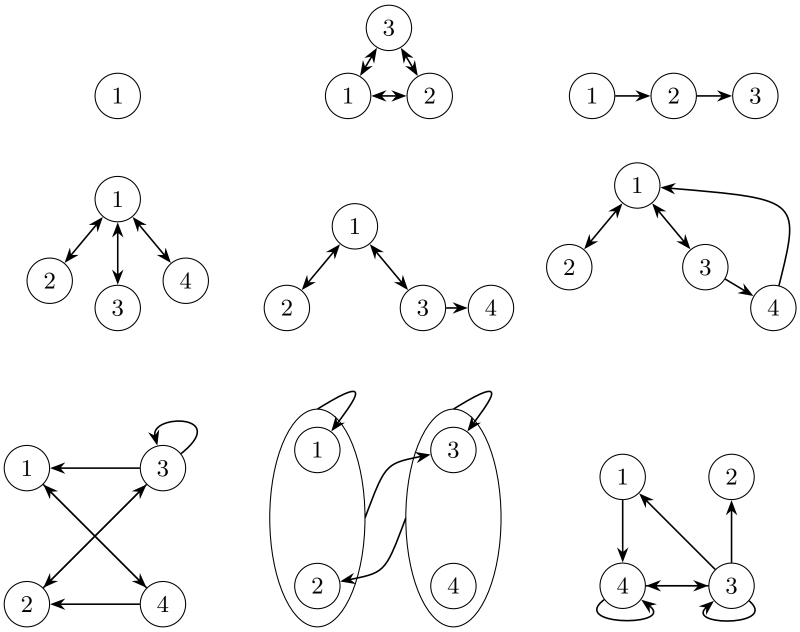

**Fig 1.** Connectivity motifs used for population coupling. In order: Null, Full, Chain, Ring, Star, Star Tail, Star Feedback Tail, Star Loop Extended, EI Extended, and an example of Erdos-Renyi connectivity motif. Arrows indicate directed connections.

#### Stochastic dynamics

Noise can enter the models in two ways: as external noisy input or as internal stochastic fluctuations. When internal noise is enabled, the builder inserts an SDE diffusion term into the first state variable of each input process block:

$$\mathrm{d}x_i = f_i \,\mathrm{d}t + \sigma_i \,\mathrm{d}W_t. \tag{104}$$

The noise amplitude $\sigma_i$ is a tunable parameter. Whenever the grammar samples a stochastic specification for a population, the corresponding diffusion term is automatically added to its state equations. External noise, however, is introduced only through the manually constructed sensory input and is not sampled by the grammar.

#### Additional details about the grammar

This appendix clarifies several aspects of the grammar that are not discussed in detail in the main manuscript.

To reduce the search space and ensure structurally sensible sampled models, we impose two context-dependent constraints during grammar sampling. First, the choice of output process depends on the number of distinct inputs received by a population. In particular, output processes that operate on a single input variable are only allowed for populations with one input process, whereas difference-based outputs are only allowed when two input processes are present. Second, when a population receives multiple inputs, the associated input processes are constrained to share the same dynamical form. This reflects a common feature of canonical neural population models, in which a given population typically integrates different incoming signals through the same synaptic or filtering mechanism.

The sampled grammar parse tree is mostly abstract: most terminal symbols specify classes of dynamical processes rather than explicit differential equations. For example, the production `InputProcess` → `Second-order` selects a second-order synaptic kernel without yet instantiating its concrete state equations. In contrast, the production `InputProcess` → `S S` recursively expands algebraic expressions and directly constructs polynomial right-hand sides during grammar expansion. The resulting formalism is therefore hybrid, combining abstract process-level model specification with direct equation generation. This enables both broad exploration of structurally constrained model families and targeted searches over explicit dynamical mechanisms.

Although these dynamical families are rarely considered within a single unified modelling space, they are not conceptually incompatible. ENEEGMA treats each population type as a modular dynamical block with well-defined inputs and outputs, enabling systematic exploration of model combinations that are uncommon in the literature yet remain mathematically coherent. This does not imply that all such hybrid models are physiologically plausible; rather, it supports structured comparison across modelling assumptions and may help identify intermediate formulations bridging established model families.

Because the recursive polynomial productions impose no a priori bound on expression length or degree, the space of admissible node models is countably infinite. The grammar can therefore generate arbitrarily complex dynamical systems while remaining within a fixed and interpretable structural template.

#### Jansen–Rit model example

To illustrate how a familiar neural mass model is generated, we use the classical Jansen–Rit (JR) cortical column as an example. Rather than hard-coding the JR equations, ENEEGMA recovers the model through a sequence of grammar expansions that progressively refines the abstract start symbol `Node` into a concrete system of coupled population dynamics.

For the JR model, the symbol `Node` is expanded into a three-population configuration together with one population-coupling function, three sensory-coupling-function slots, and one connectivity motif. This structure is obtained by expanding `Pops2` into two `Pop1` symbols, thereby fixing the total number of populations to three. Each population is then specified by the production

$$\texttt{Pop1} \rightarrow \texttt{InputProcess OutputProcess ECF ECF Stochasticity},$$

which determines how presynaptic input is transformed into postsynaptic state variables, how those states are exposed as outputs, and whether stochastic noise is present.

For the JR model, the grammar selects the second-order PSP kernel for `InputProcess` in all three populations, thereby fixing the postsynaptic dynamics to a second-order linear system. It also selects direct readout for `OutputProcess`, so that the membrane-potential-like state of each population is transmitted without additional output dynamics.

In ENEEGMA, this corresponds to instantiating the unified second-order postsynaptic potential (PSP) kernel

$$\dot{x}_1(t) = x_2(t), \tag{105}$$
$$\dot{x}_2(t) = \omega^2(g\,u(t) - x_1(t)) - 2\,\zeta\,\omega\,x_2(t), \tag{106}$$

where $x_1(t)$ denotes the lumped PSP variable and $x_2(t)$ its temporal derivative. The JR model is recovered by choosing the critically damped case $\zeta = 1$ and identifying $\omega = 1/\tau$, with $g = A$ for excitatory synapses and $g = B$ for inhibitory synapses. This yields the classical $\alpha$-kernel system

$$\dot{x}_1(t) = x_2(t), \tag{107}$$
$$\dot{x}_2(t) = A\,a\,u(t) - 2a\,x_2(t) - a^2 x_1(t), \tag{108}$$

with $a = 1/\tau$. The corresponding impulse response,

$$h(t) \propto t\,e^{-at},$$

is the characteristic $\alpha$-shaped PSP used in the original JR formulation.

To couple the populations within a node, the grammar selects the saturating sigmoid connectivity function, which implements the firing-rate nonlinearity used in the original JR model. Finally, the connectivity-motif nonterminal is resolved to a star-shaped motif, capturing the canonical three-population coupling pattern of the JR column: pyramidal-to-excitatory, excitatory-to-pyramidal, and inhibitory-to-pyramidal interactions.

Through this sequence of expansions, the grammar refines an abstract node description into a fully specified three-population neural mass model whose equations match the classical Jansen–Rit formulation up to notational differences.

### Supplementary material D: Additional results

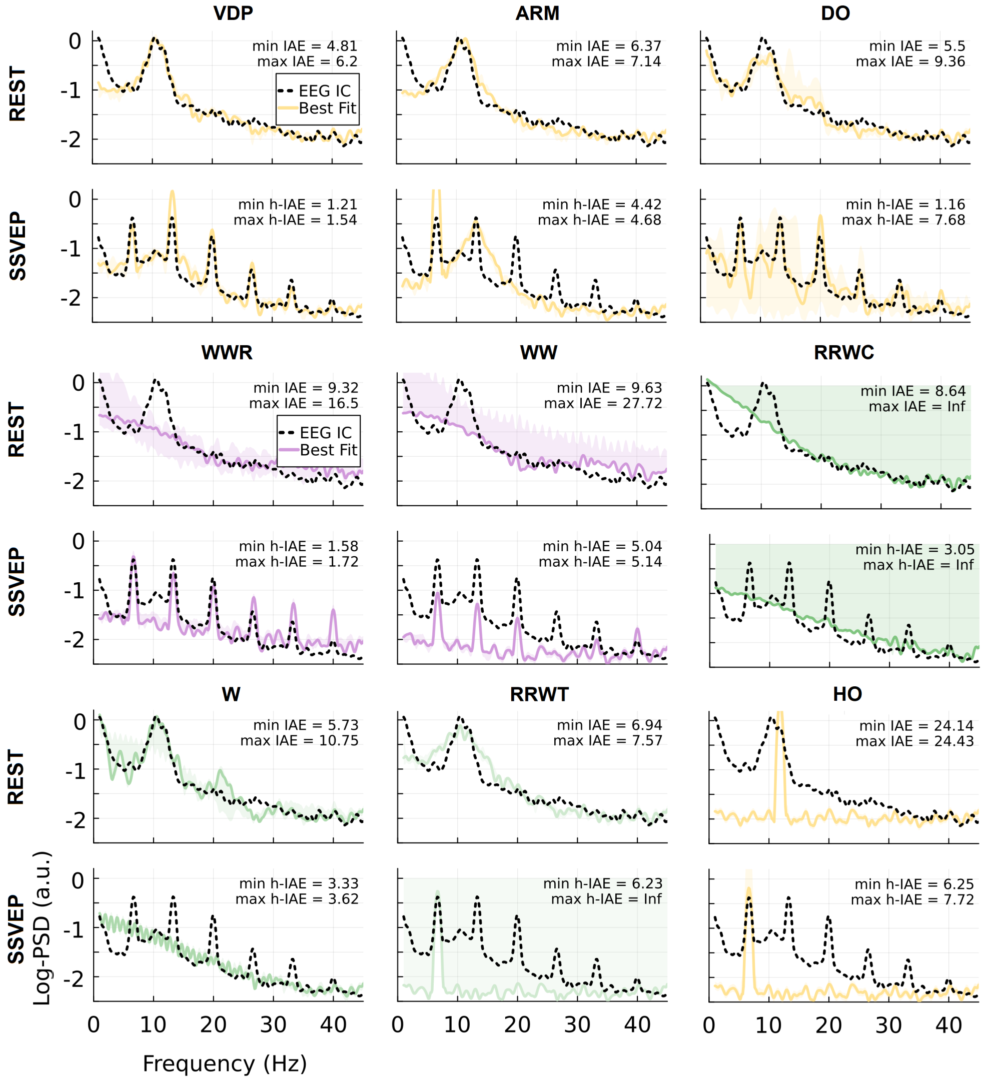


**Fig 2. Log-PSD fits of additional canonical models.** Panels show the canonical models not included in the main-text figure, evaluated for the first representative dataset under RS and SSVEP conditions. Black dashed lines denote empirical log-PSD, colored solid lines the best-fitting simulated spectra, and shaded regions the range (min–max) across five repeated simulations with different initial conditions and noise realizations. Insets report the minimum and maximum fitting error across repeated simulations (IAE for RS; h-IAE for SSVEP). As in the main text, SSVEP goodness of fit is based on harmonic-restricted error, so close agreement is expected primarily at the stimulus-locked harmonic peaks rather than across the full broadband spectrum.

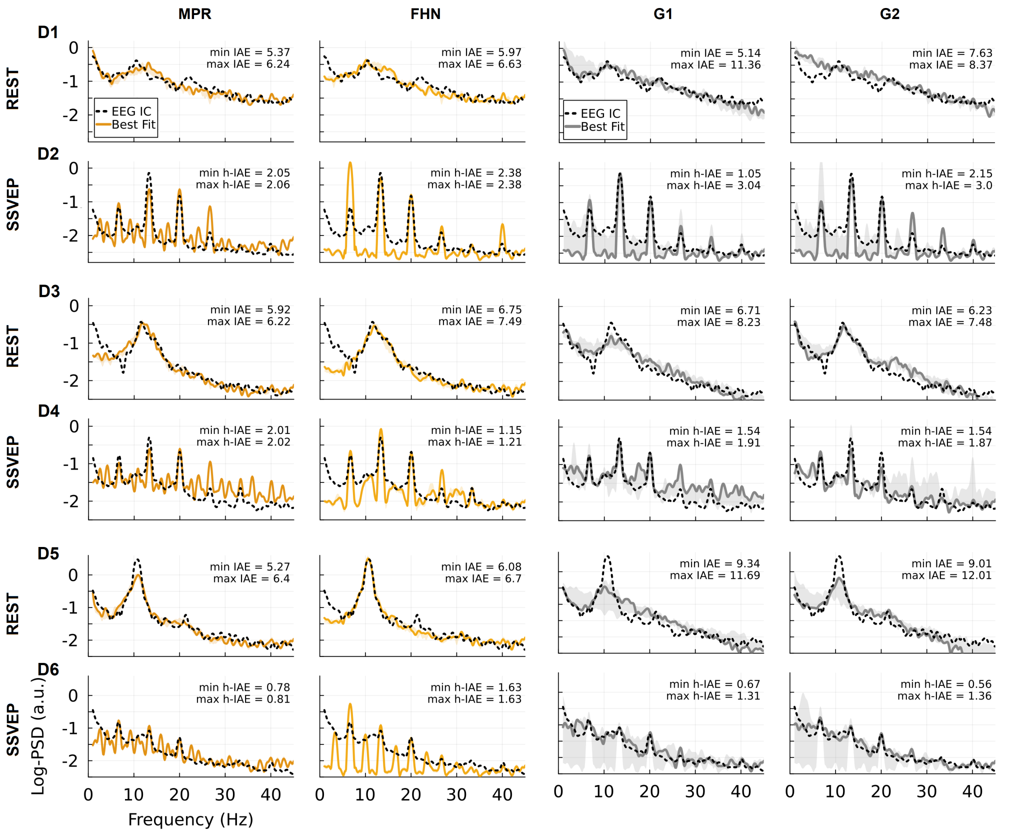

**Fig 3. Log-PSD fits of top two performing canonical and grammar-generated models across six additional datasets.** The left block shows the two top-performing canonical models (MPR and FHN), while the right block shows two top-performing grammar-generated models (G1 and G2). Each dataset is shown in a separate row pair, with RS above SSVEP. Black dashed lines denote empirical log-PSD, solid lines the best-fitting simulated spectra, and shaded regions indicate the range (min–max) across five repeated simulations with varying initial conditions and noise realizations. Insets report the minimum and maximum fitting error across repeated simulations (IAE for RS; h-IAE for SSVEP). As in the main analysis, close SSVEP agreement is expected mainly at the stimulus-locked harmonic peaks because model comparison was based on harmonic-restricted error.

#### Supplementary material E: Generated models and fitted parameters

This appendix lists the generated models discussed in the main text. For each model, we first give the reduced single-node equations in generic parametric form and then report the condition-specific fitted parameter values for the first (representative) dataset, rounded to two decimals. Because the present study considered only single-node dynamics, some distinctions present at the grammar level can disappear during reduction to the realized differential-equation system. In particular, Models G1 and G2 are symbolically identical in their reduced form, even though they arise from different grammar expansions and occupy different fitted parameter regimes across conditions.

**Model G1.**

$$\dot{x}_{11} = (c_{11} + c_{13}x_{11}x_{12} + c_{15}\,\mathrm{I}(t))\,\tau_1, \tag{109}$$

$$\dot{x}_{12} = \left(c_{12} + c_{14}x_{11}^2 + c_{16}\,\mathrm{I}(t)\right)\tau_1. \tag{110}$$

For RS, the fitted parameters for Dataset 1 were $c_{11} = 0.33$, $c_{12} = 5.73$, $c_{13} = 3.14$, $c_{14} = -0.79$, $c_{15} = -1.65$, $c_{16} = 5.75$, and $\tau_1 = 13.54$.

For SSVEP, the fitted parameters for Dataset 1 were $c_{11} = -1.30$, $c_{12} = -1.69$, $c_{13} = -1.33$, $c_{14} = 0.98$, $c_{15} = -1.74$, $c_{16} = 5.23$, and $\tau_1 = 26.93$.

**Model G2.**

$$\dot{x}_{11} = (c_{11} + c_{13}x_{11}x_{12} + c_{15}\,\mathrm{I}(t))\,\tau_1, \tag{111}$$

$$\dot{x}_{12} = \left(c_{12} + c_{14}x_{11}^2 + c_{16}\,\mathrm{I}(t)\right)\tau_1. \tag{112}$$

For RS, the fitted parameters for Dataset 1 were $c_{11} = -1.52$, $c_{12} = 6.00$, $c_{13} = 2.41$, $c_{14} = -0.35$, $c_{15} = 5.96$, $c_{16} = 3.47$, and $\tau_1 = 14.27$.

For SSVEP, the fitted parameters for Dataset 1 were $c_{11} = -0.45$, $c_{12} = 5.81$, $c_{13} = 5.54$, $c_{14} = -1.71$, $c_{15} = 6.00$, $c_{16} = 3.25$, and $\tau_1 = 7.44$.

**Model G4.**

$$\dot{x}_{11} = \left(c_{11} + c_{13}x_{12} + c_{14}x_{11} + c_{15}x_{11}^3x_{12} + c_{17}\,\mathrm{I}(t)\right)\tau_1, \tag{113}$$

$$\dot{x}_{12} = \left(c_{12} + c_{16}x_{11}^2x_{12} + c_{18}\,\mathrm{I}(t)\right)\tau_1. \tag{114}$$

For RS, the fitted parameters for Dataset 1 were $c_{11} = -1.00$, $c_{12} = -1.65$, $c_{13} = -1.51$, $c_{14} = -1.76$, $c_{15} = -1.24$, $c_{16} = -1.68$, $c_{17} = 3.30$, $c_{18} = -0.16$, and $\tau_1 = 37.92$.

For SSVEP, the fitted parameters for Dataset 1 were $c_{11} = 3.58$, $c_{12} = 3.56$, $c_{13} = -1.72$, $c_{14} = -1.01$, $c_{15} = -1.42$, $c_{16} = -1.71$, $c_{17} = 5.84$, $c_{18} = 5.95$, and $\tau_1 = 37.85$.

**Model G9.**

$$\dot{x}_{11} = \left(c_{11} + c_{15}x_{12} + c_{13}x_{11}^2 + c_{14}x_{11}^2x_{12} + c_{16}x_{11}^3 + c_{19}\,\mathrm{I}(t)\right)\tau_1, \tag{115}$$

$$\dot{x}_{12} = (c_{12} + c_{17}x_{12} + c_{18}x_{11} + c_{110}\,\mathrm{I}(t))\,\tau_1. \tag{116}$$

For RS, the fitted parameters for Dataset 1 were $c_{11} = -16.49$, $c_{12} = 22.65$, $c_{13} = 48.26$, $c_{14} = 19.81$, $c_{15} = 41.38$, $c_{16} = -17.68$, $c_{17} = 20.01$, $c_{18} = -19.43$, $c_{19} = 46.78$, $c_{110} = 26.50$, and $\tau_1 = 2.74$.

For SSVEP, the fitted parameters for Dataset 1 were $c_{11} = -1.44$, $c_{12} = -1.39$, $c_{13} = 5.82$, $c_{14} = -1.30$, $c_{15} = -1.24$, $c_{16} = -1.63$, $c_{17} = 0.90$, $c_{18} = 5.89$, $c_{19} = -0.39$, $c_{110} = 4.59$, and $\tau_1 = 24.87$.
